# Task-related, intrinsic oscillatory and aperiodic neural activity predict performance in naturalistic team-based training scenarios

**DOI:** 10.1101/2021.08.29.456571

**Authors:** Zachariah R. Cross, Alex Chatburn, Lee Melberzs, Philip Temby, Diane Pomeroy, Matthias Schlesewsky, Ina Bornkessel-Schlesewsky

## Abstract

Effective teams are essential for optimally functioning societies. However, little is known regarding the neural basis of two or more individuals engaging cooperatively in real-world tasks, such as in operational training environments. In this exploratory study, we recruited forty individuals paired as twenty dyads and recorded dual-EEG at rest and during realistic training scenarios of increasing complexity using virtual simulation systems. We estimated markers of intrinsic brain activity (i.e., individual alpha frequency and aperiodic activity), as well as task-related theta and alpha oscillations. Using nonlinear modelling and a logistic regression machine learning model, we found that resting-state EEG predicts performance and can also reliably differentiate between members within a dyad. Task-related theta and alpha activity during easy training tasks predicted later performance on complex training to a greater extent than prior behaviour. These findings complement laboratory-based research on both oscillatory and aperiodic activity in higher-order cognition and provide evidence that theta and alpha activity play a critical role in complex task performance in team environments.

## Introduction

Human and non-human primates have evolved to become highly social creatures. While social interaction is critical for the successful execution of many complex behaviours, such as teamwork, relatively little is known regarding real-world social interactions and how these are facilitated by endogenous neural activity. This, in part, is due to the difficulty in studying the neurobiological basis of social behaviour outside of the standard laboratory setting. To address this, recent work has capitalised on state-of-the-art portable electroencephalographic (EEG) devices, allowing neuroscientists to study phenomena associated with social interactions in naturalistic settings. Broadly, this work has identified several candidate neural mechanisms underlying successful social interactions within teams of two and greater, revealing alpha-mu modulations in turn-taking between a model and imitator ^1^, a negative association between single-subject alpha activity and individual-to-group alpha coherence ^2^, and that joint social action (e.g., eye contact) is associated with increased low alpha and beta synchrony ^3^. Taken together, this emerging body of work suggests a role of neural oscillations in orchestrating naturalistic social interactions ^4–6^. This extends to a larger body of work indicating that neural oscillations play a fundamental role in perceptual and higher-order information processing in the brain ^7,8^.

Neural oscillations are ubiquitous in the central nervous system, playing a pivotal role in almost all aspects of cognition. Neural oscillations during wake are typically categorised into canonical frequency bands, including theta (∼ 4 – 7 Hz) and alpha (∼ 7 – 13 Hz). Theta and alpha oscillations are two of the most widely studied neural correlates of higher-order cognition ^9^, being functionally relevant for, but not limited to, memory, attention and decision-making. From a basic neurophysiological perspective, alpha activity is thought to reflect goal-directed inhibition and disengagement of task-irrelevant brain regions ^8,10–12^, optimising the focus of attention to task-relevant information. On the other hand, theta oscillations have typically been associated with working memory and executive control functions, manifesting over fronto-central regions when recorded with scalp EEG. In operational contexts, theta activity is argued to index “mental workload”, increasing as a function of task complexity. However, from a neurobiological perspective, mental workload is difficult to define. Theoretical advances (e.g., ^13,14^) propose that theta oscillations reflect the synchronisation between the hippocampus and neocortical regions, particularly the prefrontal cortex^15^. Synchronised hippocampo-cortical theta activity may reflect a physiological mechanism for the flow of task-relevant information, with the hippocampus building successively complex representations of sensory input and the prefrontal cortex providing top-down modulations that guide behaviour and shape perception.

This framework of theta oscillations may offer a more neurobiologically plausible interpretation of theta activity reported in naturalistic contexts outside of the typical laboratory environment. For example, Diaz-Piedra and colleagues ^16^ found that theta power increased linearly with task complexity in a sample of military personnel completing simulated training exercises. Here, it was argued that an increase in theta power likely reflected aspects of problem solving and mental workload. From a physiological perspective, an increase in theta power in this context may have reflected hippocampal integration of incoming sensory input through task-relevant neocortical circuits, a process shown to depend on neural activity oscillating at the theta rhythm. However, the successful completion of real-world training, such as in a military context, likely depends on both the interaction between two or more individuals (for a detailed discussion of brain-to-brain synchrony, see ^17,18^), and individual differences in intrinsic neural activity. The interaction between such intrinsic neural activity and team performance in real-world training has not been considered in this research space to date.

Determining the nature and functional relevance of neural oscillations in higher-order cognition is complicated by the fact that individual brains oscillate around different harmonic points ^7,9^, and may process information at different speeds and timescales ^19,20^. For example, the individual alpha frequency (IAF) represents the prominent spectral peak in the alpha frequency bandwidth during eyes closed wakefulness ^21^, and is an important element in both the physiological and psychological functioning of the individual. Physiologically, IAF is proposed to determine the global speed of oscillatory activity in the brain through harmonic power laws ^7^. Cognitively, IAF may determine the rate of information processing, either through modulating temporal receptive windows for perceptual processes ^22^, or through modulating the speed and nature of the generation of internal models of the world. Regarding the former, IAF is related to the sampling rate of visual perception, such that those with a higher IAF are better able to discriminate between rapidly presented stimuli in comparison to low-IAF individuals ^22^. In terms of the latter, recent work has demonstrated that IAF modulates participants’ ability to appropriately revise their interpretations while listening to momentarily ambiguous language ^20^, and predicts the degree of sleep-based memory consolidation for various types of memory ^23,24^. In the current exploratory study, we aim to extend this nascent literature by examining intrinsic (i.e., resting-state-derived) and task-related oscillatory signatures in teams (dyads) of active-duty military personnel completing training scenarios of increasing complexity.

Military training represents an interesting context for the application of EEG to performance on demanding real-world tasks. During training, military personnel are required to perform complex tasks, often under challenging conditions (e.g., time pressure, incomplete information, simulated enemy threats, fatigue), with a strong focus on mission success, and an understanding of both task-related and intrinsic factors which may contribute towards this would represent valuable information in terms of personnel selection and in ensuring desirable operational outcomes. EEG is an ideal tool for the acquisition of neural activity under naturalistic conditions: it is relatively inexpensive and portable compared to other neuroimaging technologies (magnetic resonance imaging; magnetoencephalography), and it provides a wealth of information directly related to neural and cognitive functions (e.g., ERP components linked with attention or environmental monitoring; time-frequency representations of power changes related to memory encoding), in comparison with other physiological measurements (e.g., galvanic skin response).

Beyond studying social dynamics, naturalistic observation including EEG recording from human subjects has led to a better understanding of how the human brain processes environmental information outside of the laboratory. A recent ambulatory EEG study ^25^ demonstrates that the mismatch negativity (MMN) and P300 in an auditory oddball paradigm are reduced in environments outside laboratory settings, potentially due to increased demands for (complex) environmental monitoring. This highlights a need for *in situ* monitoring of complex behavioural tasks to fully understand the neural correlates of human behaviour therein, and thereby predict outcomes. This extends beyond task-locked brain activity, and can include less obvious performance-related factors, such as intrinsic measures of neural activity obtained from periods of resting wakefulness.

As with the IAF, aperiodic elements of the human EEG (broadly, non-oscillatory activity thought to reflect the excitation/inhibition balance of neural networks) can be used to predict brain states (i.e., sleep/wake) as well as clinical status (e.g., schizophrenia, attention-deficit hyperactivity disorder; ^26^). Resting-state-derived oscillatory and aperiodic elements of the EEG change in response to environmental conditions. For example, individuals isolated for 120 days under space analogue conditions showed a broad decrease in aperiodic factors in the EEG (including a flattening of 1/ƒ slope), and cessation of isolation lead to a temporary increase in IAF ^26^. It is, however, currently unknown how both task-and resting-state-derived oscillatory and aperiodic activity predicts performance in complex, naturalistic environments in groups of individuals.

Using state-of-the-art portable EEG, the current exploratory study aimed to identify electrical brain activity associated with team performance for military dyads completing simulated training under naturalistic conditions. Here, military teams completed simulated tank gunnery training or simulated ground-based air defence (GBAD) training tasks using in-service simulation systems. The tank gunnery training involved military personnel completing virtual battlefield scenarios in dyads, comprised of a commander and gunner. Each dyad was required to successfully communicate to make optimal decisions and reach target objectives (i.e., detect, engage, and destroy enemy vehicles) across three scenarios of increasing complexity. Similarly, the GBAD training involved military personnel completing virtual air defence scenarios in teams of two (commander and gunner) across three scenarios of increasing complexity.

We aimed to address the following research questions: (1) are there EEG-based correlates of performance on simulated training tasks in a military context, and do these markers track performance over time and increasing task difficulty? (2) In the case of an association between EEG-based markers and task performance, are these markers more precise predictors of performance than established behavioural markers (e.g., subjective assessment by trained professionals)? (3) Are there resting-state derived measures of brain activity that predict task performance, and does this differ depending on task demands and roles (e.g., gunner and commander)?

## Results

### Descriptive statistics

Overall, commanders reported having been in their current role (quantified as months) longer than gunners in the Armoured (commanders, *M* = 59.7, *SD* = 48.74; gunners, *M* = 28.7, *SD* = 20.65) and GBAD (commanders, *M* = 53.6, *SD* = 29.52; gunners, *M* = 19.2, *SD* = 9.30) groups. Commanders also reported longer military service experience than gunners in the Armoured (commanders, *M* = 111.5, *SD* = 49.41; gunners, *M* = 67.0, *SD* = 15.23) and GBAD (commanders, *M* = 84.5, *SD* = 44.27; gunners, *M* = 27.3, *SD* = 9.35; see Figures 1A and 1B) samples. For dyad performance in the Armoured and GBAD samples, see Figures 1C and 1D, respectively.

**Figure 1.**
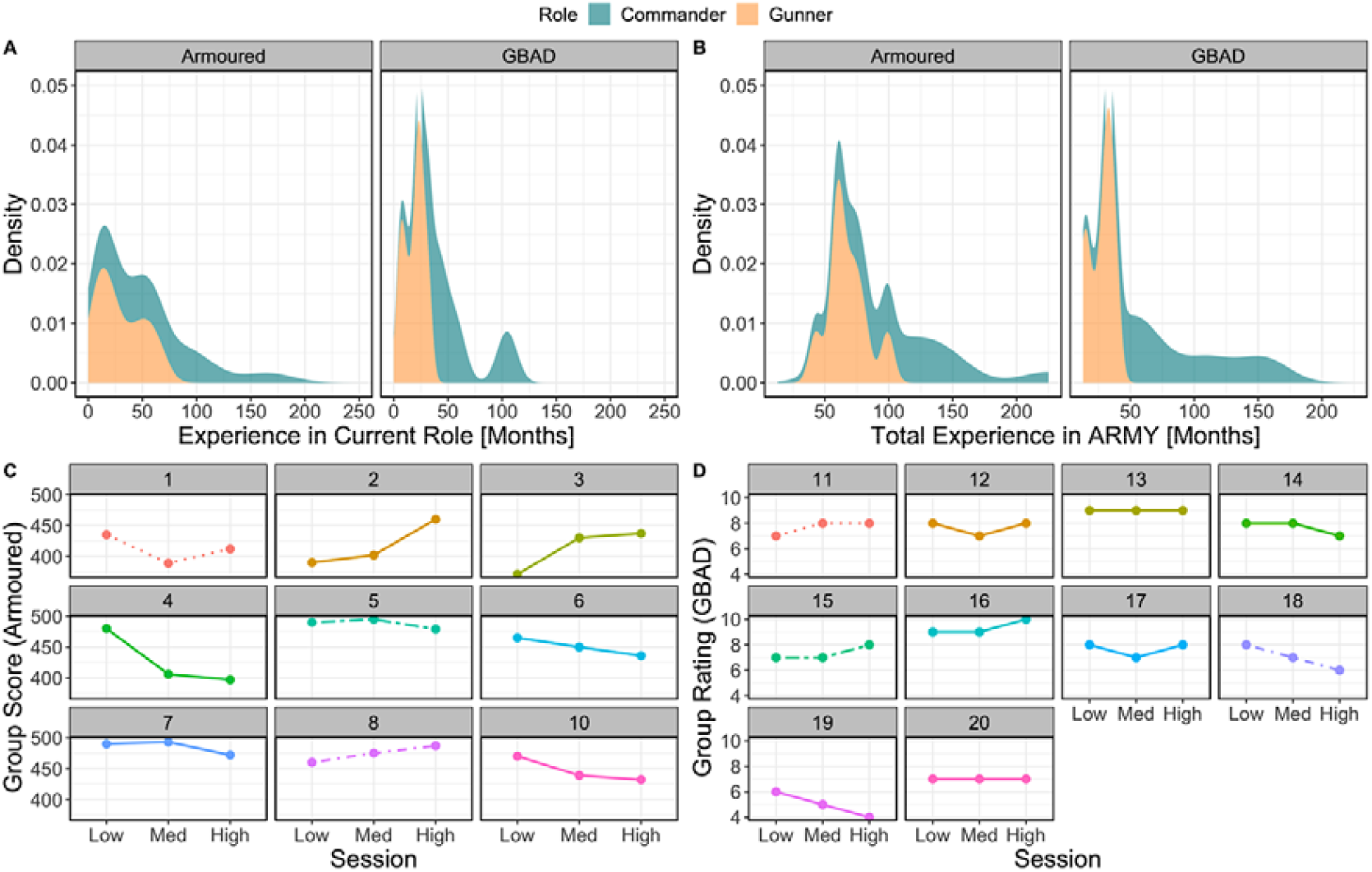
Summary of time served in the military and dyadic behavioural performance. (A) Summary of time (in months) in current role between gunners (orange) and commanders (blue) at Armoured (left) and GBAD (right), while (B) illustrates the same for total time in military service (months). (C) Performance scores for each dyad during the Armoured tank simulation task. Each facet represents a dyad (i.e., gunner and commander). Performance is represented on the y-axis (higher values indicate better performance), while session (one = low, two = med, three = high) is represented on the x-axis. (D) Subjective rating scores for performance for each dyad during the GBAD dome simulation task. Each facet represents a dyad (i.e., gunner and commander). Performance is represented on the y-axis (higher values indicate better performance), while training scenario (easy, moderate difficult) is represented on the x-axis.

### Training complexity is associated with task-related theta and alpha power

Here, we ran linear mixed-effects models to determine whether theta and alpha power change across scenarios one to three (easy to difficult), between gunners and commanders and between GBAD and Armoured groups. The results from these models are illustrated in Figure 2A and 2B. For the theta model, there was a Role × Session × Study interaction (χ^2^(2) = 88.72, *p* < .001). As is clear from Figure 2A, theta power estimates between commanders and gunners in both groups (GBAD and Armoured) were roughly equivalent during the easy session but diverged during the moderate and highly difficult scenarios. Here, theta power increased the most for commanders, particularly GBAD, while theta power estimates decreased for the Armoured gunners.

**Figure 2.**
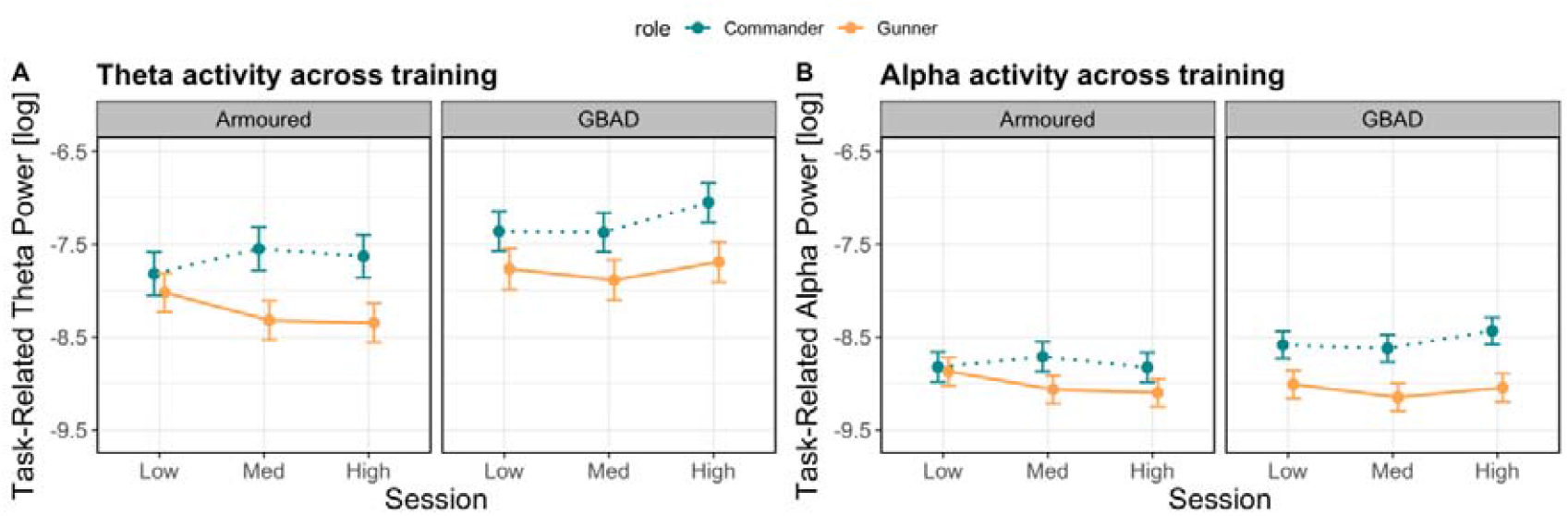
Modelled effects of task-related neural activity. (A) Theta power (y-axis; higher values indicate increased power) across scenario one (Low), two (Med) and three (High; x-axis) between Armoured (left) and GBAD (right). Estimates from commanders are represented by the dashed blue line, while estimates from gunners are represented by the solid orange line. Error bars represent the 83% confidence interval. (B) illustrates the same as (A) but for alpha power.

The alpha model also revealed a significant Role × Session × Study interaction (χ^2^(2) = 38.49, *p* < .001). Alpha power estimates were overall higher for commanders relative to the GBAD gunners. For Armoured, alpha activity decreased from the easy to difficult session for gunners but increased from the easy to moderately difficult session before tapering off thereafter for Commanders. Taken together, these results indicate that modulations in theta and alpha power reflect changes in task complexity, but also depend on the role of an individual and the specific training regime.

### Theta and alpha activity predict behavioural performance across task complexity

Given the clear difference in task-related theta and alpha activity across the easy, moderate, and difficult scenarios between the gunners and commanders, we performed analyses to test whether theta and alpha activity under different levels of task complexity predict performance. Linear mixed-effects modelling revealed modulations in GBAD dyad performance ratings as a function of task-related theta and alpha power (for a visualisation of all modelled effects, see Figure 3). For the theta model, there was a significant Power × Role × Session interaction (χ (2) = 68.11, *p* < .001): while theta activity did not predict performance ratings in the easy and moderately difficult scenarios for commanders, higher theta power was associated with better performance for gunners in both scenarios (Figure 3A). This pattern reverses in the difficult scenario, where an increase in theta power was associated with improved performance for commanders, but not gunners.

**Figure 3.**
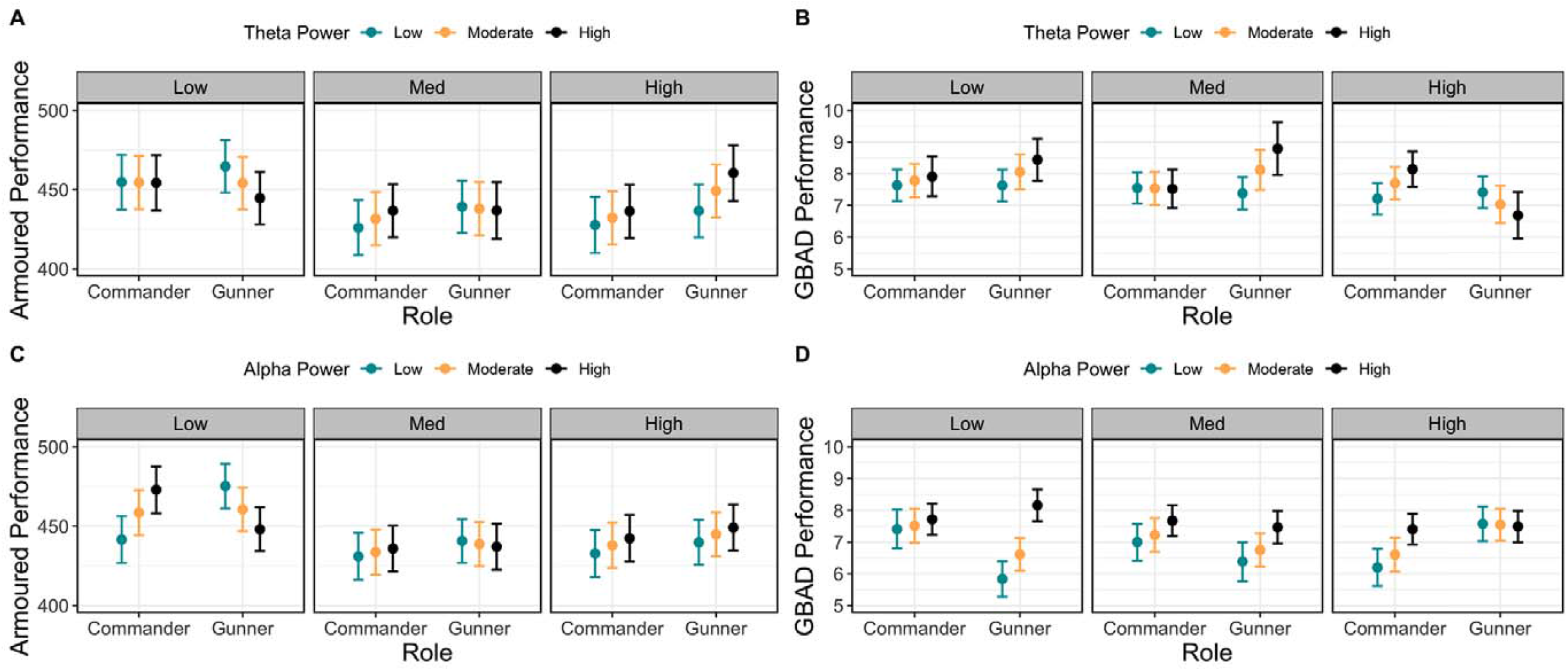
Modelled effects of task-related theta and alpha activity on behavioural performance. (A) Relationship between performance ratings (y-axis; higher values indicate better performance), role (commander, gunner; x-axis), session (easy = left; medium = middle; high = right), and task-related theta power (blue = low power; yellow = moderate power; black = high power). Bars represent the 83% confidence interval. (C) represents the same as (A) but for task-related alpha power, while (B) and (D) represent the same but for performance on the tank simulator at Armoured. Here, performance is represented on the y-axis, with higher values indicating better performance. Note that power values (theta and alpha power) were discretised into low, moderate, and high values based on the first quartile, median and third quartile values for the purposes of plotting but were entered into the model as continuous predictors.

The alpha model also revealed a significant Power × Role × Session interaction (χ^2^ (2) = 39.33, *p* < .001; see Figure 3C). Here, an increase in alpha power was associated with higher performance ratings for the gunners in the easy and moderately difficult scenarios, but not in the difficult scenario. By contrast, while alpha power showed no association with performance for commanders in the easy session, an increase in alpha power was associated with improved performance ratings in the moderate and difficult scenarios.

For performance in the tank gunnery task, there was a significant Power × Role × Session interaction for the theta band (χ^2^(2) = 14.60, *p* < .001). While theta power was not predictive of performance for commanders on any of the scenarios, lower theta power was associated with better performance on the easy scenario for gunners, while this pattern reverses in the difficult scenario (Figure 3B). The alpha model also showed a significant Power × Role × Session interaction (χ^2^(2) = 44.73, *p* < .001). Here, in the easy scenario, an increase in alpha power predicted improved performance for commanders, while the inverse is observed for gunners (Figure 3D). This pattern of results weakens in the moderately difficult session, and in the difficult scenario, an increase in alpha power was predictive of improved performance in both gunners and commanders, with the magnitude of this effect being greater for gunners.

### High and low performing dyads show differing task-related activity across training difficulty

In order to examine team-based performance, we categorised dyads into either low or high performers based on whether the dyad scored below or above mean group performance across each session, respectively. We then ran a linear mixed-effects regression using the following formula:

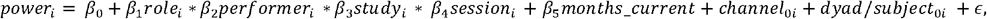

Here, *power* is task-related oscillatory power in the theta or alpha bands, *role* encodes gunner and commander, *performer* refers to low or high performing dyads, *study* is Armoured or GBAD studies, and *session* encodes low, medium and high difficulty training scenarios. *Months current* was entered as a covariate in order to account for months served in current role, while *channel* and *subject* were specified as random effects, with *subject* nested under *dyad*. ε refers to a Gaussian-distributed error term. Critically, categorising dyads into low and high performers enabled us to somewhat overcome the non-randomisation of session difficulty, while also examining team performance.

The model predicting theta power revealed a significant Role × Performer × Study × Session interaction (χ^2^(2) = 193.80, *p* < .001). This interaction is resolved in Figure 4A, where, for the Armoured study, gunners in low performing dyads in the low difficulty session had reduced theta power, while commanders had increased theta power. For high performing dyads, theta power was equivalent between gunners and commanders. This pattern is similar in the highly difficult scenario, suggesting that gunners and commanders in high performing dyads had similar profiles of theta activity during both easy and difficult training scenarios in the Armoured context.

**Figure 4.**
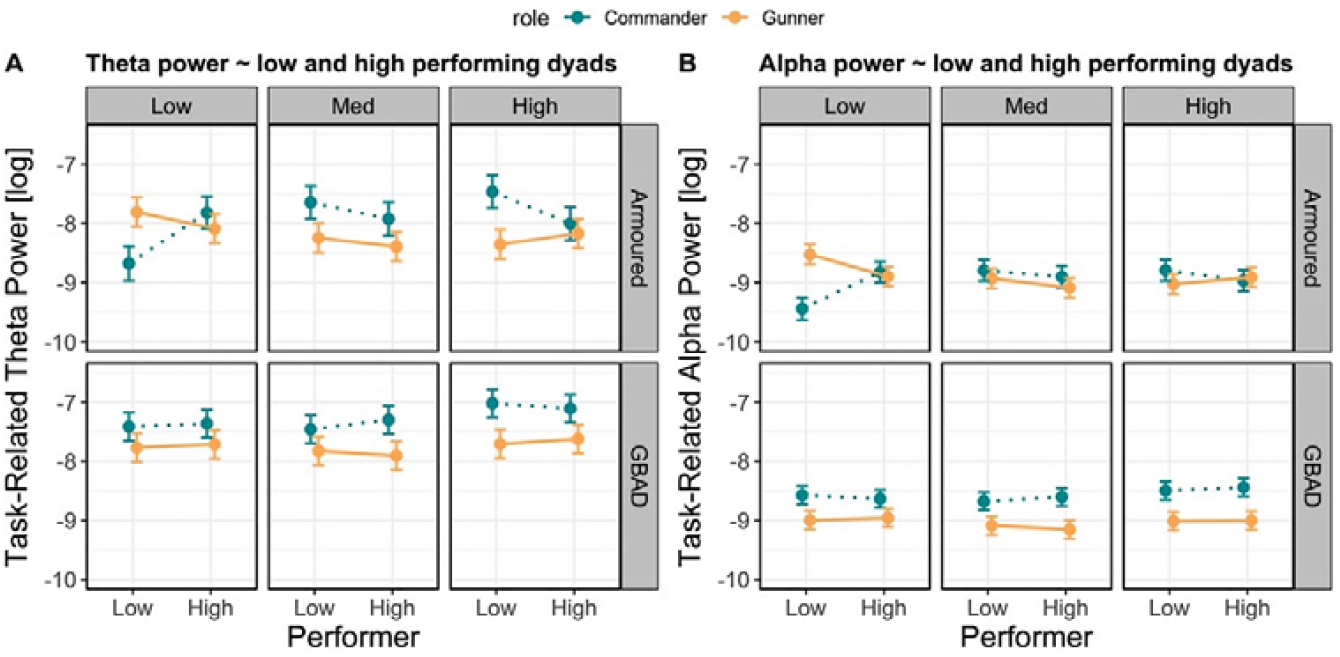
Modelled effects of task-related theta and alpha activity between low and high performing dyads. (A) Relationship between task-related theta power (y-axis; higher values indicate greater power), performer (x-axis; left = low performing dyad, right = high performing dyad), session (easy = left; medium = middle; high = right), and study (top row = Armoured; bottom row = GBAD). Estimates from commanders are represented by the dashed blue line, while estimates from gunners are represented by the solid orange line. Error bars represent the 83% confidence interval. (B) represents the same as (A) but for task-related alpha power.

The model examining alpha power also revealed a significant Role × Performer × Study × Session interaction (χ^2^(2) = 250.25, *p* < .001). As shown in Figure 4B, during the low difficult session in the Armoured study, high performing dyads had similar alpha profiles, while low performing dyads reveal differential patterns of alpha activity. Taken together, the theta and alpha models suggest that high performing dyads in the Armoured study have similar profiles of oscillatory activity, while gunners and commanders in low performing dyads have differential patterns of oscillatory activity.

### Task performance under complex conditions is predicted by intrinsic and prior task-related neural activity

Next, we focussed on whether we can predict performance on the most difficult scenario (session three) based on both task performance and EEG indices from the easiest scenario (the first session) in both groups (i.e., Armoured and GBAD). Here we used generalised additive mixed models (GAMMS) to model non-linear relationships between our predictors (e.g., task-related neural activity) and performance, given that many biological phenomena do not fit simple linear models (e.g., ^27,28^).

The analyses for Armoured revealed that performance on the easy scenario was not predictive of performance on the difficult scenario (*β* = 0.06, *R*^*2*^ = -.20, *p*_adj_ = .71; Figure 5A); however, task-related alpha power explained approximately 26.6% of the variance in performance (Figure 5B). This effect was largest for the commander, with performance from session three showing a curvilinear relationship with alpha power from session one (edf = 2.46, *F* = 4.22, *p*_adj_ < .001). Task-related theta power from scenario one also showed a curvilinear relationship with performance on scenarios three, explaining approximately 42.1% of the variance in performance. Here, this effect was largest for the commanders (edf = 2.51, *F* = 5.61, *p*_adj_ < .001).

**Figure 5.**
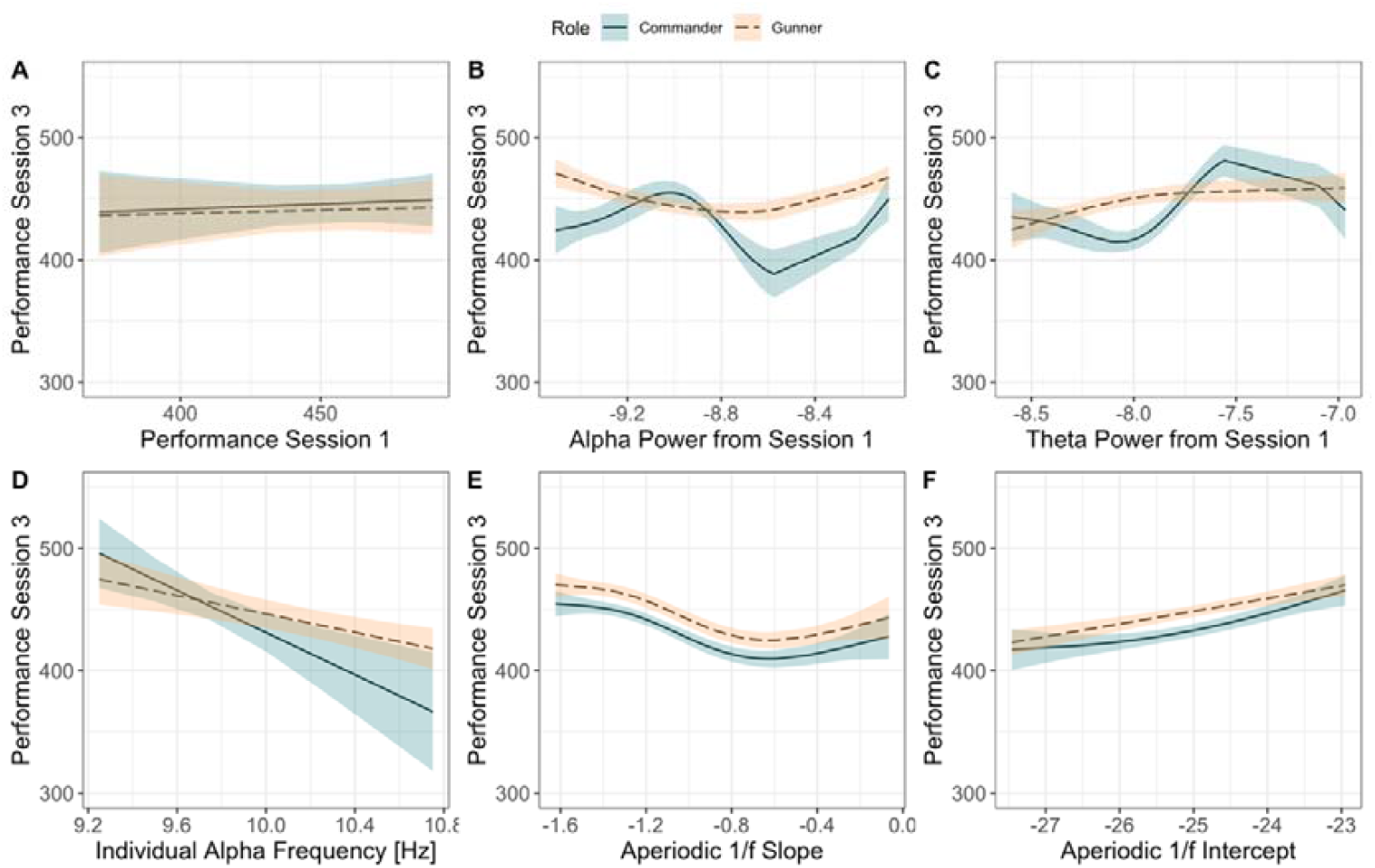
Modelled effects of behavioural, task-related, and resting-state-derived EEG activity on behavioural ratings from session three for the simulated tank task at Armoured (most difficult scenario; represented on the y-axis, with higher values indicating better performance).

Resting-state-derived 1/ƒ slope (Figure 5E) explained performance over and above that of the performance ratings, explaining 42.7% of the variance (edf = 2.98, *F* = 15.61, *p*_adj_ < .001), an effect that was not modulated by role. Similarly, the 1/ƒ intercept (Figure 5E) explained 24.1% of the variance in performance from session three, with both the gunner and commander showing a linear increase in performance with a decrease in the 1/ƒ intercept (edf = 9.38, *F* = 3.81, *p*_adj_ < .001). Finally, resting-state-derived IAF explained approximately 45% of the variance in performance from session three (*β* = -61.89, *p*_adj_ = .008; Figure 5D). While this effect did not statistically vary by role, the slope was steeper for commanders than gunners, with performance decreasing as IAF increased.

For GBAD performance, task-related alpha activity in the first scenario explained the largest amount of variance in performance during the final scenario (*R*^*2*^ = .28), which is visualised in Figure 6B. Here, alpha power from session one shows a U-shaped effect on performance during session three for the commander (edf = 3.54, *F* = 11.05, *p*_adj_ < .001), but a positive linear relationship with performance for the gunner. By contrast, behavioural ratings from scenario one (easiest condition) explained 18% of the variance in performance (*β* = 0.83, *R*^*2*^ = .18, *p*_adj_ = .03; Figure 6A); however, this effect did not vary by role.

**Figure 6.**
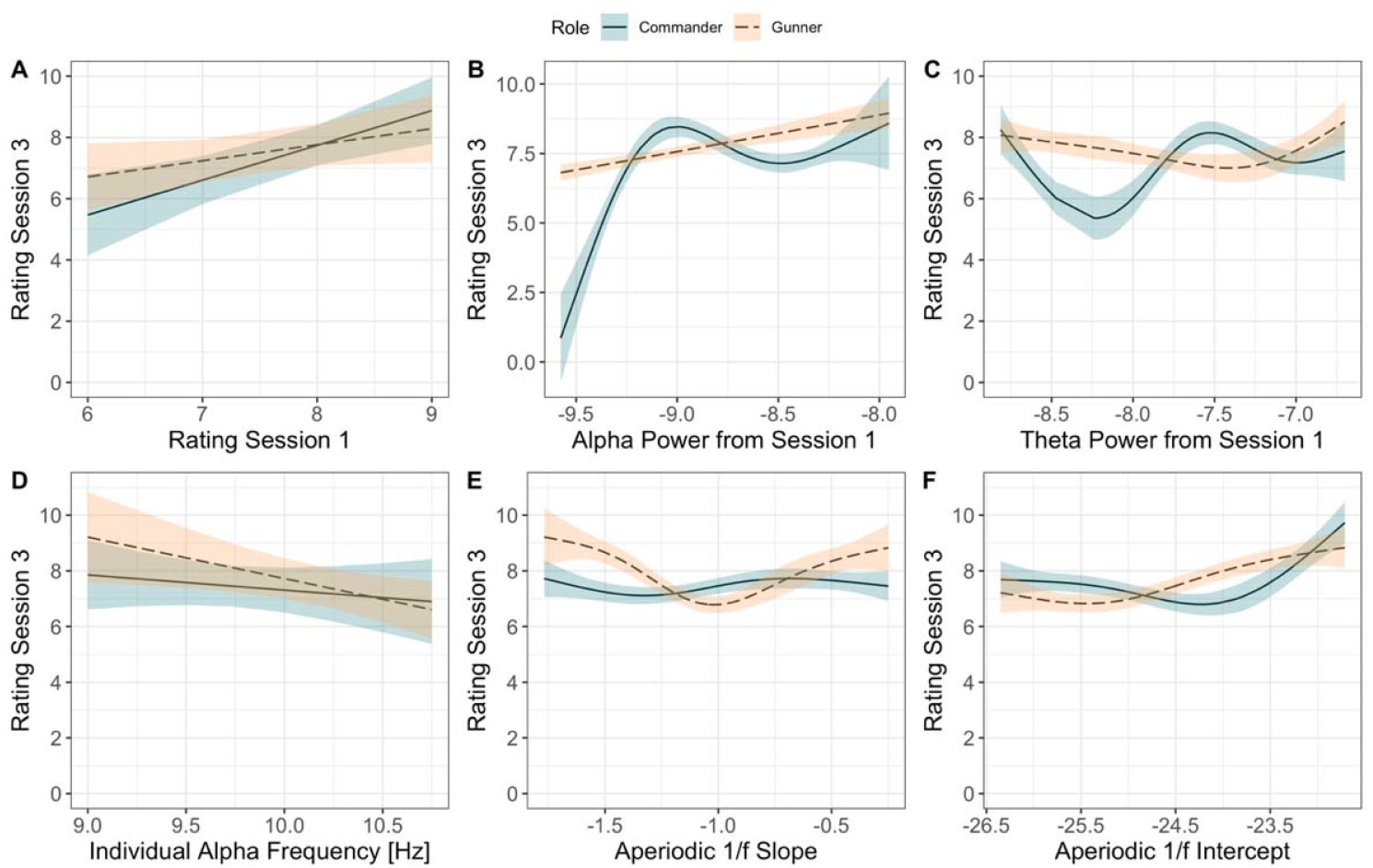
Modelled effects of behavioural, task-related, and resting-state-derived EEG activity on performance ratings from session three for the GBAD task (most difficult training scenario; represented on the y-axis, with higher values indicating better performance).

Resting-state-derived 1/ƒ intercept (Figure 6F) explained 17.4% of the variance. This effect was strongest for the commander, with performance lower when the intercept was higher, before showing a slight U-shaped increase when the intercept decreased (edf = 2.03, *F* = 4.28, *p*_adj_ < .001). For the 1/ƒ slope (Figure 6E), there was a U-shaped relationship with performance for the gunner (edf = 2.21, *F* = 6.12, *p*_adj_ < .001), explaining approximately 14.6% of the variance in performance.

Task-related theta (Figure 6C) power was also predictive of behavioural performance on the most difficult scenario three (*R*^*2*^ = .17), with the model revealing a strong non-linear effect of theta power on performance for both the commander (edf = 2.70, *F* = 4.70, *p*_adj_ < .001) and gunner (edf = 1.64, *F* = 1.38, *p*_adj_ = .06; uncorrected *p* = .03); however, the relationship between theta power and performance for gunners was no longer significant after Holm-Bonferroni adjustment. The IAF explained the least amount of variance in performance (*β* = -1.01, *R*^*2*^ = .01, *p*_adj_ = .13; Figure 6D).

### Predicting role identity based on intrinsic neural activity

As an exploratory analysis, we examined whether there were differences in three metrics between gunners and commanders: resting-state derived IAF, and the 1/ƒ slope and 1/ƒ intercept. In order to control for differences in age between the gunners and commanders which may drive IAF-related effects in the logistic regression machine learning model, we ran a simple linear regression predicting IAF from Age, revealing that age in years was significantly negatively associated with IAF (*β* = -0.03, *R2* = .13, *p* = .02). From this, we extracted the residuals from the IAF ∼ Age linear regression (using the *broom* package v.0.7.5), with the residuals reflecting the unexplained variance between IAF and Age. The residuals were then entered into the logistic regression machine learning model, replacing IAF. Differences between gunners and commanders on these metrics are visualised in Figure 7, which show differences in IAF, the 1/ƒ intercept, age and the residuals from the IAF ∼ Age regression between gunners and commanders. In order to determine whether we could classify role (gunner, commander) based on resting-state EEG metrics, we constructed a logistic regression machine learning model using the *R* package *tidymodels* v.0.1.2. Data were separated into a training and test set, retaining 75% of the data for training and the rest for testing. We then created bootstrapped resamples of the training data to evaluate our logistic regression model, which took the following form:

**Figure 7.**
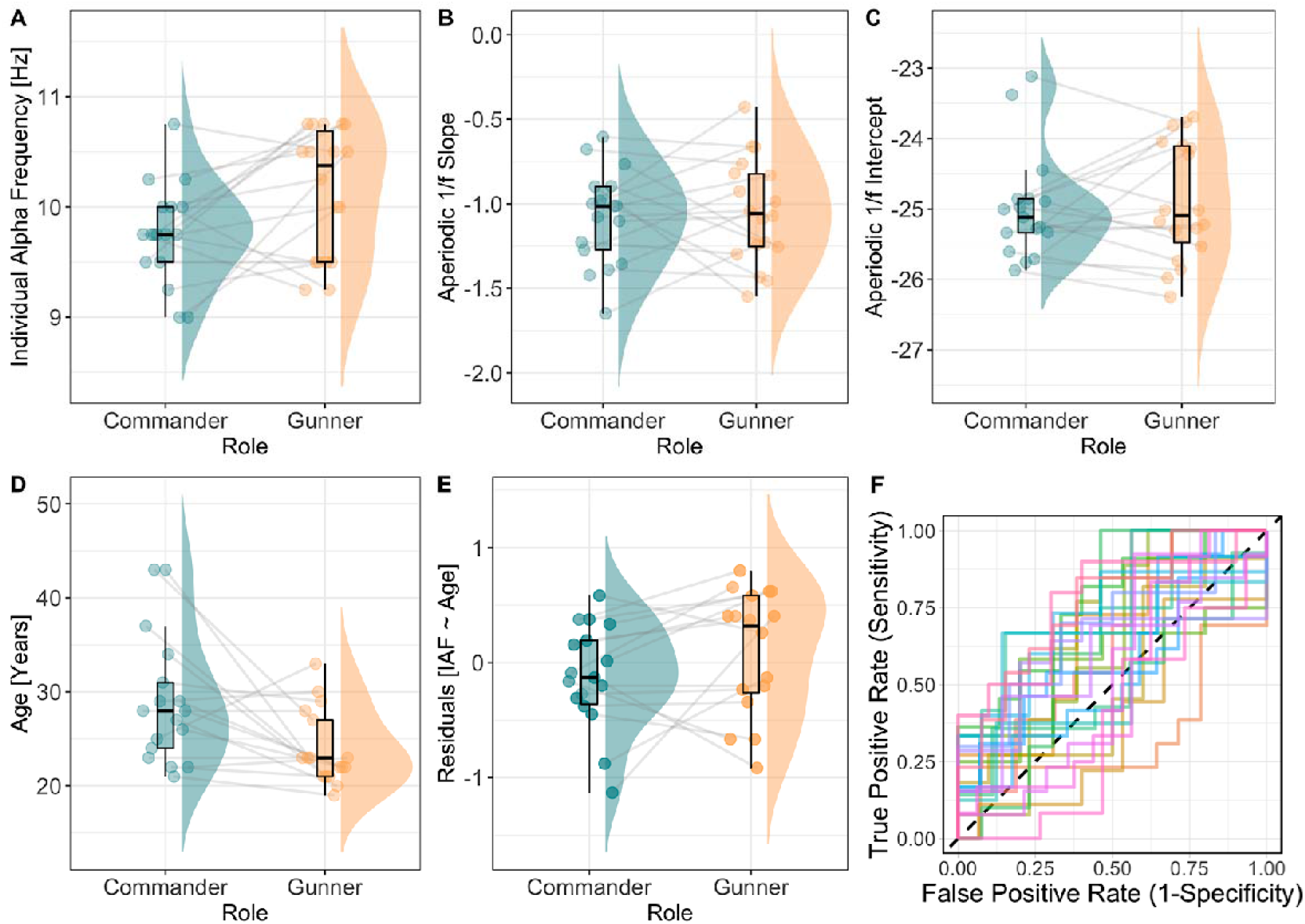
Boxplots illustrating difference in resting-state EEG between gunners and commanders. (A) IAF is represented on the y-axis, with higher values indicating a higher IAF. (B) The 1/ƒ slope is represented on the y-axis, with higher values reflecting shallower slopes. (C) The 1/ƒ intercept is represented on the y-axis, with higher values indicating a higher intercept. (D) Age (years) is represented on the y-axis, with higher values representing older age. (E) Residuals from the IAF ∼ Age regression are represented on the y-axis, with positive values representing residuals above the regression line, negative values below the regression line, and values of zero indicating data points that are intersected by the regression line. Data points reflect individual participants, with joined data points between gunners and commanders representing specific pairs (e.g., dyads). (F) Region over the curve plot representing the logistic machine learning model. Each line represents one of the 25 bootstrapped resamples of the training data.

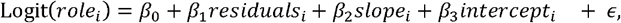

Here, *role* refers to the binary outcome of gunner or commander, *residuals* is the residuals from the IAF ∼ Age regression, *slope* represents the aperiodic 1/ƒ slope, and *intercept* is the aperiodic 1/ƒ intercept. The model performed well on the test data with a region under the curve estimate of .67. To examine the individual predictive capacity of each factor, we exponentiated the coefficients to produce odds ratios (OR), revealing that the residuals from the IAF ∼ Age regression was the strongest predictor of role. For every one unit increase in residuals, an individual had 3.70 times higher odds of being a gunner than a commander (*se* = .60, *p* = .03). The 1/ƒ slope (OR = 3.84, *se* = 1.54, *p* = .39) and the 1/ƒ intercept (OR = 1.97, *se* = .43, *p* = .12) were not predictive of role. To determine whether the predictive power of IAF on role was also not confounded by experience, we conducted two linear regressions predicting IAF from length of military service and time in current role. Neither model was predictive of IAF (service time: *β* = -0.002, *R2* = .06, *p* = .36; role experience: *β* = -0.002, *R2* = .04, *p* = .66).

## Discussion

Here, we have recorded brain activity from dyads of defence personnel performing military training exercises and used these data to gain insights into how intrinsic neural (IAF and 1/ƒ slope and intercept) and task-related oscillatory EEG activity (alpha and theta power) relate to task performance. Broadly, our results demonstrate: (1) differential patterns of task-related alpha and theta power as a function of individual role and task complexity; (2) that these task-related oscillatory indices are related to operational outcomes, and; (3) that intrinsic (resting-state-derived) oscillatory and aperiodic metrics are predictive of behavioural outcomes on the training task, even demonstrating predictive capacity above and beyond that of task performance from less complex training scenarios. These data provide important insights for understanding the ways in which the human brain functions in complex team environments and provide insights which could potentially be applied towards team selection and the training of personnel for specific roles and tasks.

It is presently assumed that successful team performance (defined in broad behavioural terms) must rely on shared representation of task structure and rules for interaction and performance (for a detailed discussion, see ^29^). In terms of underlying neural mechanisms, it is assumed that interbrain synchrony or other neural activity relates directly to success, and this is driven either by shared or overlapping bottom-up, sensory processing between team members, or the shared deployment of cognitive resources, such as attention to relevant factors and operations ^2^. It has, however, proved difficult to disentangle these two explanatory mechanisms.

Our results regarding the functional relevance of task-related oscillatory activity, showing broadly similar patterns of alpha and theta albeit differences in task complexity, suggest that high performance in team settings may be measured through the EEG, potentially tagging shared, task-relevant deployment of attentional and memory processes ^7,9^; however, future work should directly examine whether measures of inter-brain synchrony between gunners and commanders predict behavioural performance. This interpretation is supported by the finding that task-related alpha power from session one can predict task performance on session three, explaining more variance than task performance on session one. This finding suggests that EEG-based measures can provide additional insight regarding predicted performance on a complex, naturalistic training scenario over and above performance on a more basic scenario. It is important to note, however, that there were differences in oscillatory activity depending on role, with gunners showing a linear relationship between alpha power on a simple scenario and performance on a complex scenario and commanders showing a curvilinear one. This difference may relate to the distinct attentional strategies used by gunners and commanders: the narrow focus of attention required for optimal performance in gunners (requiring inhibition of all irrelevant stimuli) and the broader attentional focus and situational awareness required by commanders (too much inhibition could be detrimental to performance, as important environmental stimuli could fail to be incorporated into the broader situational view) may be reflected by the linear and U-shaped relationship between alpha power and performance, respectively. The linear relationship for gunners describing the necessity to maintain object tracking despite environmental distractors, and the U-shaped relationship for commanders demonstrates a necessary mix of inhibition and situational awareness. It must be noted, though, that such a clean division of cognitive labour likely does not exist in the brain, and other processes are likely also at play. Future work should employ complementary measures to EEG that track cognitive processing and team performance. For example, concurrent eye-tracking and EEG, in addition to the analysis of vocal communication between gunners and commanders (i.e., via voice recordings), would help to further decompose attentional modulations across session difficulty and successful team performance, respectively. A finer temporal resolution of the processing of stimuli via the use of triggers to critical events would also offer a more fine-grained insight into the neurophysiological mechanisms underlying training in the current context. For example, one might expect reductions in attention-related event-related potentials (e.g., P300) as session difficulty increases, serving as a proxy for attentional adaptation in complex team-based training. Further, it is interesting to note that performance on the Armoured study was more consistent with traditional laboratory-based experiments. This may be due to the objective simulator generated metrics of performance (compared to the GBAD study which relied on expert ratings). From this perspective, the use of objective simulator-based metrics, in conjunction with eye-tracking and voice recording analyses and stimulus triggers, may provide a deeper insight into how the human brain performs under complex team-based military training.

Individual differences in the EEG – which underlie key differences in cognitive-behavioural performance – should also be considered. Similar to the discussion around the neural sequelae of group performance described above, perspectives differ as to whether IAF may influence individual-level performance through the modulation of temporal perceptive windows ^22,30^, or via the gating of information processing, resulting in differences in the generation and manipulation of internal, generative models of the world and predicted, upcoming sensory stimuli ^20^, or elements of both. While our study is not specifically constructed to test these ideas, our results can still be considered in light of the debate around the mechanisms through which IAF influences cognition and behaviour. Using machine learning techniques, we were able to predict the role of a participant based on their IAF. In a somewhat similar vein to the explanation for task-related alpha power above, the differences in IAF may reflect different aptitudes for information processing. A faster sampling of the sensory environment, as reflected by a higher IAF, may be advantageous for gunners, as a higher environmental sampling rate is associated with a higher resolution of visual information and thus possibly with more accurate sensorimotor performance. By contrast, a lower IAF, and thus a slower sampling of the sensory environment may be critical for commanders, as a slower sampling rate would allow for better integration of information across broader timescales and from multiple sources of information. This finding provides an initial indication that resting-state EEG metrics, as stable markers of cognitive capacity, might be useful in selecting personnel for suitable roles, such as early identification of individual aptitude for progressing to the role of commander in crews such as those examined in the present study. Thus, individual differences in the EEG are predictive of performance in real-world tasks and may inform selection criteria for these. These concepts are reinforced by considering other EEG metrics, such as the 1/ƒ slope and intercept.

While IAF is relatively well understood in terms of functional relevance, aperiodic factors in the EEG are comparatively less understood in the broader cognitive neuroscience literature. One prominent hypothesis is that the aperiodic slope indexes the inhibition/excitation balance in the brain ^31,32^, and our results indicate that both the slope and intercept are highly predictive of behavioural outcomes (i.e., task performance), outperforming even prior behaviour as a marker of subsequent performance in complex, naturalistic scenarios. For example, we found that performance became worse alongside a flattening of the 1/ƒ slope. This finding is in line with established literature, which has generally noted performance improvements on complex tasks with steeper resting-state 1/ƒ slopes ^31,33,34^. The 1/ƒ slope is hypothesised to reflect the excitation–inhibition balance, or the ratio between excitatory and inhibitory cell activity in a neural population. Interestingly, a shift toward either excitation or inhibition has been proposed to impair function ^32^, which may offer a functional interpretation for our complex non-linear relationship between the 1/ƒ slope and performance of both the Armoured (Figure 5E) and GBAD (Figure 6E) dyads. From this perspective, there may be an “optimal” state of aperiodic activity, reflecting intrinsic, inter-individual differences in performance capabilities. An interesting avenue for future research may thus involve determining whether the 1/ƒ slope can be (temporarily) modified either through cognitive training interventions or non-invasive brain stimulation methods (e.g., transcranial direct current stimulation) as a way of optimising performance outcomes in complex naturalistic settings. Similarly, a lower 1/ƒ intercept was strongly related to performance, potentially due to the proposed function of 1/ƒ intercept as a broad marker of population neural firing, and thus overall network communication. It is curious that we found an inverse relationship between 1/ƒ intercept and outcomes given this, but this finding may reflect habituation or expertise on the training task as our subjects were all experienced soldiers, and the task was part of their regular occupational duties.

Naturalistic studies carry with them increased noise in comparison to traditional, laboratory-based experimentation. This extra noise (both as artefact such as sweat and sway/movement, and the effects of noise of other latent environmental stimuli) is unavoidable and is, to some extent, a side-effect of the increased ecological validity of the approach. It is important to note, however, that in both studies reported herein, participants completed all tasks in climate-controlled conditions, helping to mitigate sweat-related modulations in electrode impedance. Increased noise necessitates increased precision in theoretical models and measurement of constructs of interest. There is presently no adequate theoretical model to explain differences in naturalistic versus non-naturalistic task processing in the brain. Further, some theorists have argued that the determination of causality in studies involving two or more individuals is impossible without the use of multi-subject brain stimulation techniques ^35^. As such, future work should employ brain stimulation methods to help determine the causal relationship between brain activity from groups of individuals and behaviour, an approach which would be complemented by strong methodological and experimental design ^35,36^.

Taken together, in this exploratory study we have shown how task-related and resting-state-derived oscillatory and aperiodic factors in the EEG can be used to gain insight into human performance in complex, naturalistic, team-based settings. We have demonstrated that measures of aperiodic and resting state EEG can be used to predict behavioural performance, sometimes with greater predictive capacity than previous behaviour. We have also shown that it is possible to differentiate between members within dyads, based on resting state measures. Results of our analyses can provide information to facilitate optimal performance in occupational settings and to improve our understanding of the mechanics which allow the human being to collaborate and to succeed in real-world tasks and environments.

## Method

### Participants

Forty adults currently serving in the Australian Defence Force who were drawn from armoured and artillery regiments took part in the study (see Table 1 for a summary of participant demographic information). All participants reported no history of psychiatric, neurological, cognitive or language disorders, normal or corrected-to-normal vision, right-handedness and no use of medications that may affect EEG. Ethics approval was obtained from the University of South Australia’s Human Research Ethics Committee and the Defence Science and Technology Group (DSTG) Low-Risk Ethics Committee prior to study commencement

**Table 1.** Sample size and demographic information from the armoured and artillery samples.

|  | Armoured |  | Artillery/GBAD |  |
| --- | --- | --- | --- | --- |
|  | Commander | Gunner | Commander | Gunner |
| *n* | 10 | 10 | 10 | 10 |
| Age | 29.20 (6.33) | 26.60 (3.43) | 28.00 (6.96) | 21.22 (1.20) |
| Sex (male) | 10 | 10 | 8 | 9 |

*Note.* Standard deviation for age is presented in parentheses.

### Simulation training tasks

For all training tasks, participants used in-service simulators developed for the purposes of military training and undertook standard training scenarios. In the Armoured context, participants completed a simulated tank task, which simulated the dynamics of the vehicle over several terrains. The simulation environment presented participants with multi-modal cues (i.e., visual, auditory, motion), including audio cuing from a console operator, as well as vehicle and ammunition sounds. The simulation included three scenarios, each increasing in complexity. Across each scenario, commanders and gunners were required to work together to engage with simulated enemy combatants, distinguishing between targets to be engaged and fired upon and to avoid engaging with non-targets. Performance metrics on the tank gunnery simulation training task were taken from performance analysis provided by the simulator, reflecting the time taken to identify and successfully engage with enemies, as well as rounds fired and accuracy of the engagement. Final scores for each training scenario were aggregated measures of engagements, with a maximum score of 500.

For the GBAD training scenarios, participants also worked in pairs (commander and gunner) to coordinate their actions against simulated enemy air threats. The commanders and gunners were provided with visual, audio and motion cues in an immersive virtual reality environment. The dome screen for the virtual environment has scenarios projected from 27 projectors arranged in three rows of nine, all mounted from a central chandelier structure. The total projected image covers 270 horizontal degrees of the 12-metre dome. In this environment, commanders were required to track the position of potential enemy air threats using a hand-held geoplot device that provided coordinates of the potential threats. Once identified, the commanders directed the gunners to where the potential enemy threat was located. If a threat, gunners engaged with the enemy using a simulated RBS70 air defence launcher. Two raters (who were experienced training instructors) reached a consensus on the performance of the dyads using a 10-point rating scale, with 0 denoting poor performance and 10 denoting excellent performance. The experts also rated the individual performance of the commanders and gunners using the same scale, yielding individual-and dyadic-level performance metrics. For a visualisation of the GBAD training scenario, see Figure 8.

**Figure 8.**
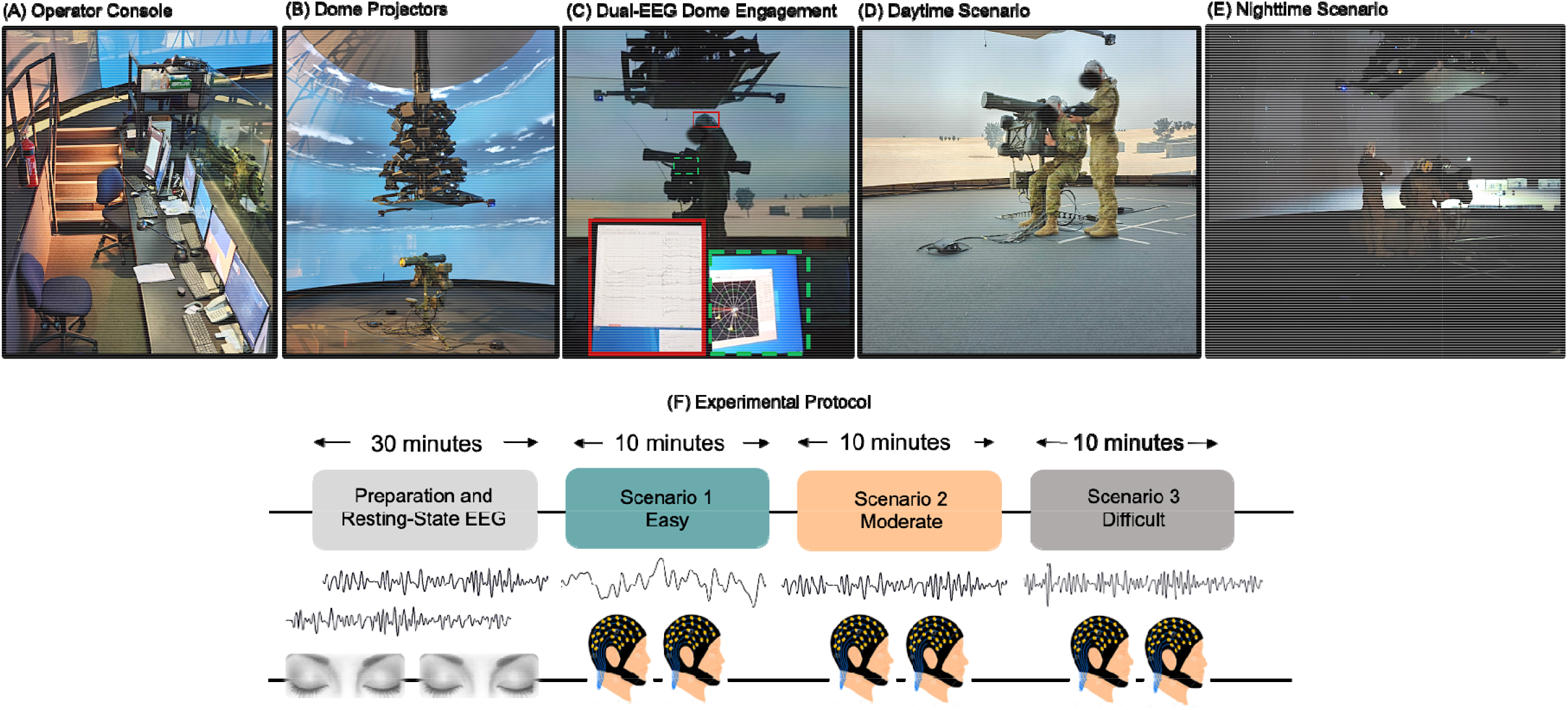
Illustration of GBAD simulation environment and experimental protocol. (A) Console operator quarters. Here, the scenarios were controlled, and each dyad was rated by two expert trainers. (B) Twenty-seven mounted projectors that displayed the training scenarios. (C) Example of EEG recording (coded in red solid square) system and the information displayed to the commander (coded in green dashed square). (D) and (E) illustrate examples of different testing scenario conditions in the ground to air simulation training. (D) Daytime scenario utilised in scenarios one (easy) and two (moderate). (E) Night-time condition utilised in scenario three (difficult). (F) Schematic of the testing protocol used for both samples. Thirty-minutes was allocated for EEG cap preparation for the gunner and commander and two minutes of resting-state EEG. Each dyad (i.e., pair of gunner and commander) then completed training sessions of increasing complexity, with each training session lasting for approximately ten minutes.

### Protocol

Prospective participants first completed an initial screening form to determine their eligibility. If eligible, participants provided written consent to participate. The gunner and commander were both fitted with EEG (comprising a single dyad) and completed two minutes of resting state EEG recording with eyes closed prior to the first training scenario.

During the tank gunnery task, the gunner and commander operated as a pair, working together to quickly navigate, spot and react to targets that they found in the virtual battlespace. Both visual and auditory information were provided to participants to facilitate successful task completion. For the virtual air defence task, participants similarly worked in pairs to coordinate their actions against simulated enemy air threats. Participants completed seven 10-minute training sessions, with each session increasing in difficulty. We were interested in tracking changes in performance and EEG as a function of task complexity, and collected EEG data from three of these sessions to capture performance on baseline/easy, moderate, and difficult tasks. EEG was continuously recorded throughout these sessions. EEG was only recorded from three sessions due to logistical reasons (e.g., battery life of the portable EEG system). Refer to Figure 8 for a diagram of the experimental protocol.

## Data Analysis

### EEG recording and pre-processing

EEG was recorded simultaneously from each dyad (i.e., gunner and commander) at rest and throughout the tank and ground-to-air simulation training using two LiveAmps (Brain Products GmbH, Gilching, Germany) with 32 active Ag/AgCl electrodes mounted in elastic caps (Brain Cap, Brain Products GmbH, Gilching, Germany). Electrode placement followed the 10/20 system. Eye movements and blinks were monitored with frontal electrodes (Fp1/Fp2). All channels were amplified using a LiveAmp amplifier (LiveAmp 32, Brain Products, GmbH) at 500 Hz. All EEG pre-processing and analysis was performed using MNE-Python ^37^. Raw data were band-pass filtered from 0.1 to 30 Hz (zero-phase, hamming windowed finite impulse response [FIR filter; 16,501 sample filter length; 0.1–7.5 Hz transition bandwidth]). Data were then re-referenced to the average of TP9 and TP10. Artefacts were corrected using Infomax Independent Component Analysis ^38^ and the Autoreject package ^39^ (for modelling of electrooculography-related independent components across session, see the supplementary material). EEG segments were also dropped when they exceeded a 150 μV peak-to-peak amplitude criterion or were identified as containing recordings from flat channels (i.e., < 5 μV). For task-related recordings, the continuous EEG signal for each training session was segmented into fixed length epochs of 30s using the function mne.make_fixed_length_epochs.

### Resting-state neural activity

Individual alpha frequency (IAF) estimates were derived from two minutes of resting-state EEG recordings taken prior to the first training session. IAF estimates were taken from occipital-parietal electrodes (P3/P4/O1/O2/P7/P8). This IAF estimation routine uses a Savitzky-Golay filter (frame length = 11 frequency bins, polynomial degree = 5) to smooth the power spectral density (PSD). It then searches the first derivative of the smoothed PSD for evidence of peak activity within a defined frequency interval (here, 7 – 13 Hz). For full methods, see^21^.

To extract the aperiodic components (1/ƒ slope and intercept) from resting-state EEG recordings, we used the irregular-resampling auto-spectral analysis method (IRASA v1.0; ^40^) implemented in the YASA toolbox in MNE-Python. IRASA isolates the aperiodic (random fractal) component of neural time series data via a process that involves resampling the signal at multiple non-integer factors *h* and their reciprocals 1/*h*. This resampling procedure systematically shifts narrowband peaks away from their original location along the frequency spectrum, averaging the spectral densities of the resampled series attenuates peak components, while preserving the 1/ƒ distribution of the fractal component. The exponent summarising the slope of aperiodic spectral activity is then calculated by fitting a linear regression to the estimated fractal component in log-log space, an example of which is provided in Figure 9. For a full mathematical description of IRASA, see ^28,40^.

**Figure 9.**
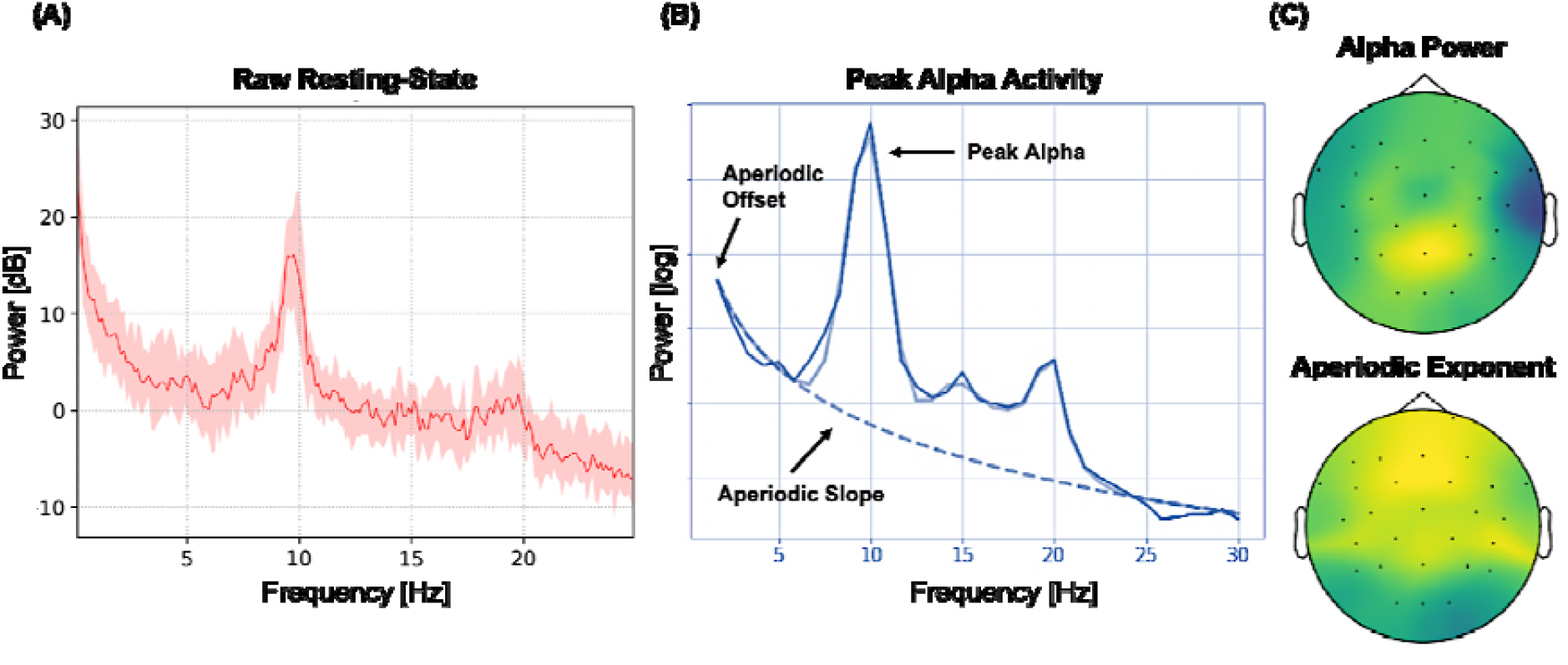
Exemplar power spectral density (PSD) estimates of fractal (aperiodic 1/ƒ) and oscillatory activity from a single participant. (A) PSD plot from two minutes of an eyes-closed resting state period. Power (dB) is presented on the y-axis, while frequency (Hz) is on the x-axis. The solid line represents the mean power across all channels, while the shaded area represents the standard error of the mean. (B) Separation of oscillatory (dark solid blue and solid light blue lines) and aperiodic (dashed blue line) components ^31^. Power (log) is on the y-axis, while frequency (Hz) is on the x-axis. Note the clear alpha oscillatory peak at approximately 10 Hz. (C) Top: topographical distribution of alpha power within the peak alpha range as shown in (B). Bottom: topographical distribution of the aperiodic exponent. Warmer colours denote greater alpha power and a higher aperiodic exponent (i.e., steeper slope).

### Task-related time-frequency analysis

Task-related time frequency analyses were performed in MNE-Python using a family of complex Morlet wavelets via the function tfr_morlet. Individualised (based on IAF) theta (∼ 3 – 7 Hz) and alpha (∼ 8 – 13 Hz) bands were analysed using wavelet cycles for each 30 second epoch. From this, we derived theta and alpha power estimates from each scenario, study (Armoured, GBAD), role (commander, gunner) and channel (Fz, F3, F7, FT9, FC5, FC1, C3, CP5, CP1, Pz, P3, P7, O1, Oz, O2, P4, P8, CP6, CP2, Cz, C4, FT10, FC6, FC2, F4). For a visualisation of the pre-processing and time-frequency analyses, see Figure 10.

**Figure 10.**
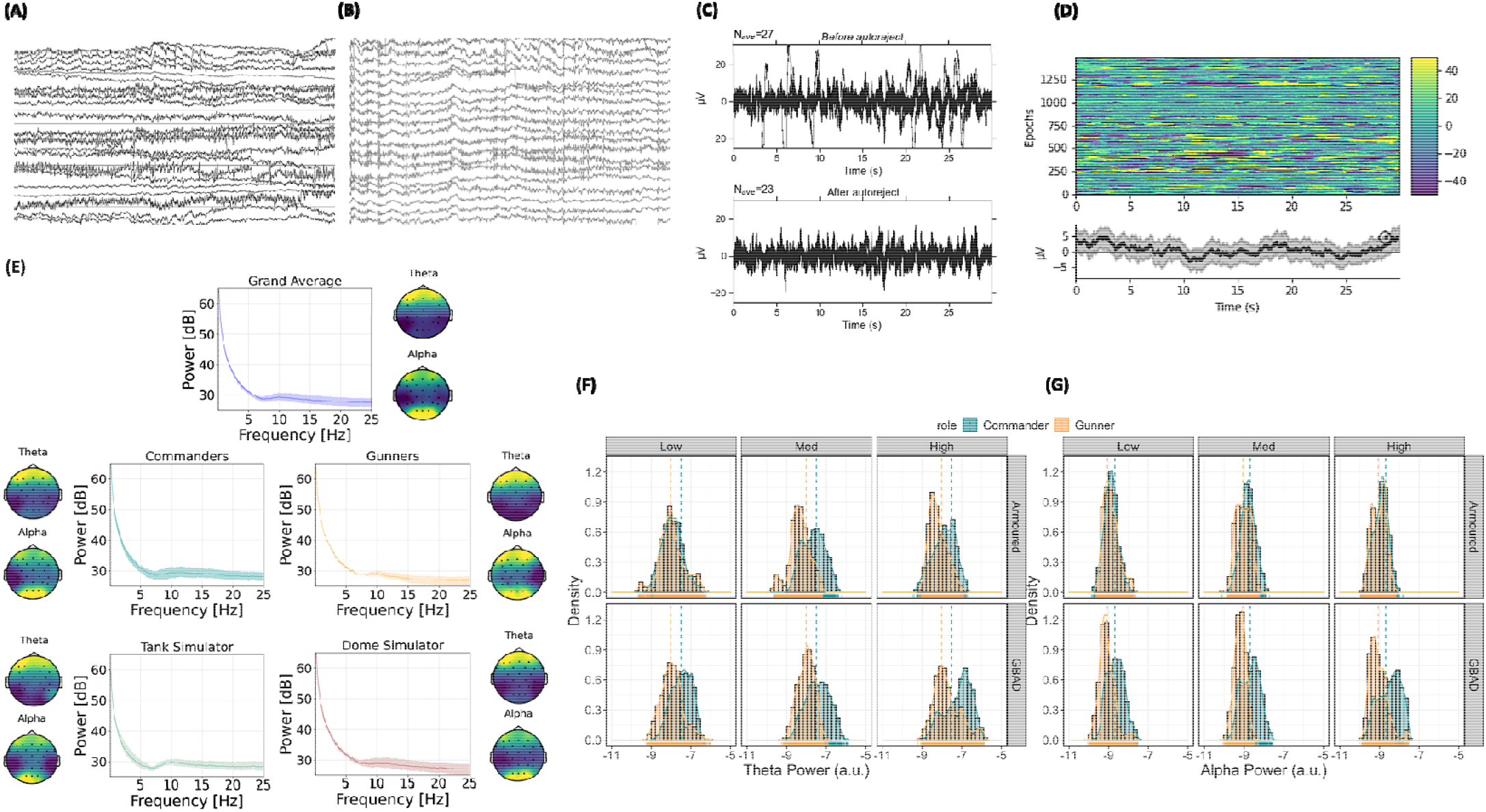
Visualisation of the pre-processing and time-frequency decomposition of task-related theta and alpha activity from the real-world training scenarios. (A) and (B) illustrate the raw and filtered and re-referenced EEG trace for a single subject, respectively, while (C) represents the change in the EEG signal before and after applying Autoreject ^39^. (D) illustrates amplitude (μV) across epochs (y-axis) and time (x-axis) for every subject at channel Cz (top figure). The bottom figure illustrates the average amplitude (y-axis; μV) across time (x-axis) for all epochs. (E) shows the power spectral density across each subject (Grand Average), commanders, gunners, tank simulator (armoured) and dome simulator (GBAD), with corresponding topographical distributions of theta and alpha power. (F) and (G) illustrate raw distributions of theta and alpha power estimates, respectively, from Armoured (top row) and GBAD (bottom row), facetted from training sessions one (easy; left), two (medium; middle) and three (high; right), for both commanders and gunners.

### Statistical analysis

Statistical analyses were conducted using *R* version 3.6.2 (R Core Team, 2020) with packages *tidyverse* v.1.3.0 ^41^, *car* v.3.0.8 ^42^, *effects* v.4.1.4 ^43^, *mgcv* v.1.8-31 ^44^, *mgcViz* v.0.1.4 ^45^, *lme4* v.1.1.26 ^46^, and *tidymodels* v.0.1.2 ^47^. Plots were created in *R* using *ggplot2* v.3.3.0 ^48^, while *lmerOut* v.0.5 was used to produce model output tables, and *ggeffects* v.1.0.2 ^49^ was used to extract modelled effects for visualisation.

#### Linear mixed-effects models

Linear mixed-effects models were used to examine differences in theta and alpha power during each of the training scenarios between gunners and commanders, and GBAD and Armoured (tank training). Role (gunner, commander), Task Complexity (easy, medium, high difficulty) and Study (Armoured or GBAD) were specified as fixed effects with full interactions. Subject ID and electrode were modelled as random effects on the intercept, while dyad was modelled as a random slope nested under Subject ID. Power (*power*) was specified as the outcome. Months experience in participants’ current role (commander, gunner) was also modelled as a covariate to control for any influence of experience on task-related EEG and behavioural performance. Type II Wald χ^2^-tests were used to provide *p*-value estimates, while an 83% confidence interval (CI) threshold was adopted for visualisations, which corresponds to the 5% significance level with non-overlapping estimates ^50,51^. Study (Armoured, GBAD) and Role (Gunner, Commander) were entered as unordered factors using sum-to-zero contrast coding (reference category coded -1), while Session (easy, medium, high difficulty) was specified as an ordered factor. Note that when contrast coding is explicitly described, the need for post-hoc testing is eliminated (for a detailed discussion of contrast coding in linear mixed-effects regressions, please see ^58^). For generalized additive and linear regression models, the Holm–Bonferroni method was used to correct for multiple comparisons.

#### Generalised additive mixed-effects models

Generalised additive mixed-effects models (GAMMS) are extensions of linear mixed-effects models, containing both fixed and random effects, as well as being capable of replacing linear predictors with a smooth function ^52–54^. This smooth function allows for the modelling of non-linear relationships, such as, modulations in behaviour as a function of neural activity (see ^27,28^ for similar approaches). Here, we used GAMMS to examine how behavioural performance is predicted by (non-linear) modulations in task-related theta and alpha activity, as well as resting-state-derived IAF and aperiodic metrics.

Behavioural performance was modelled as a function of session and task-related theta and alpha activity. We ran separate models for each frequency band (theta, alpha) and resting-state-derived EEG metrics (IAF, 1/ƒ slope, 1/ƒ intercept) to determine whether variations in these predictors modulate behavioural performance scores. Performance during session three (the most complex scenario) were modelled as a function of performance during session one (the least complex scenario) or task-related EEG (theta, alpha power) or resting-state-derived neural activity (IAF, aperiodic slope or intercept) and role (commander, gunner). A random factor smooth of channel (i.e., non-linear equivalent of a random effect in a linear mixed-effects regression) was also included to account for topographic differences in task-and resting-state-related oscillatory activity.

GAMMs were estimated using the *bam*() function of the *R* package *mgcv* ^55^. Models were fit using the Fast REML method and with tensor product interaction smooths. Tensor product smooth interactions enabled us to examine main effects and interactions in an ANOVA-style format, including estimated degrees of freedom (edf; ^56^). All tensor product smooths were fit using low rank thin plate regression splines as their basis function ^54,57^. Theta and alpha power estimates were also log_10_ transformed prior to inclusion in all models, and to isolate outliers, we used Tukey’s method, which identifies outliers as exceeding ± 1.5 × inter-quartile range.

## Acknowledgements

The authors would like to thank MAJ Tim Pexton for his invaluable role as an Army liaison during data acquisition as well as the military personnel who participated in this study. We are also grateful to Sophie Jano and Scott Coussens for their help with data acquisition, and Colin O’Donnell and Kelly Witt for their support with the simulation systems used in this study. Thank you also to Andrew Corcoran for valuable feedback on an earlier version of this manuscript, and to LTCOL Bevan McDonald and Dr David Crone for their support to the study, especially their input during the conceptualisation and planning stages.

