## supplementary material for "Task-related, intrinsic oscillatory and aperiodic neural activity predict performance in naturalistic team-based training scenarios"

Table S1: Summary of model produced by the call `lmer(formula = power ~ role * session * study + Months_Current + (1 | ch_name) + (1 | dyad/subj), data = tf_power_theta, REML = TRUE, control = lmerControl(optimizer = "bobyqa", calc.derivs = TRUE))`
Linear mixed model fit by REML. t-tests use Satterthwaite's method

REML criterion at convergence: 502

Scaled residuals:

| Min | 1Q | Median | 3Q | Max |
|---|---|---|---|---|
| -4.01 | -0.63 | -0.02 | 0.57 | 4.16 |

Random effects:

| Groups | Term | Std.Dev. |
|---|---|---|
| subj:dyad | (Intercept) | 0.42042 |
| ch_name | (Intercept) | 0.10895 |
| dyad | (Intercept) | 0.20491 |
| Residual | | 0.24842 |

Number of obs: 2658, groups: subj:dyad, 39; ch$_name, 25; dyad, 20.$

Fixed effects:

| | Estimate | Std. Error | df | t value | Pr(>\|t\|) | |
|---|---|---|---|---|---|---|
| (Intercept) | −7.7 | 0.13 | 34 | −58 | 1.7e−35 | *** |
| role[Commander] | 0.27 | 0.08 | 21 | 3.4 | 0.0029 | ** |
| session[Low] | −0.0073 | 0.0071 | 2.6e+03 | −1 | 0.31 | |
| session[Med] | −0.047 | 0.0071 | 2.6e+03 | −6.7 | 2.4e−11 | *** |
| study[Armoured] | −0.21 | 0.083 | 17 | −2.6 | 0.019 | * |
| Months$_Current$ | −0.0011 | 0.0025 | 31 | −0.44 | 0.66 | |
| role[Commander]:session[Low] | −0.12 | 0.0071 | 2.6e+03 | −17 | 3.1e−59 | *** |
| role[Commander]:session[Med] | 0.05 | 0.0071 | 2.6e+03 | 7.1 | 1.5e−12 | *** |
| role[Commander]:study[Armoured] | 0.011 | 0.068 | 16 | 0.16 | 0.88 | |
| session[Low]:study[Armoured] | 0.035 | 0.0071 | 2.6e+03 | 4.9 | 9.9e−07 | *** |
| session[Med]:study[Armoured] | 0.061 | 0.0071 | 2.6e+03 | 8.6 | 1.4e−17 | *** |
| role[Commander]:session[Low]:study[Armoured] | −0.062 | 0.0071 | 2.6e+03 | −8.6 | 1e−17 | *** |
| role[Commander]:session[Med]:study[Armoured] | 0.053 | 0.0071 | 2.6e+03 | 7.5 | 7.1e−14 | *** |

Table S2: Summary of model produced by the call `lmer(formula = power ~ role * session * study + Months_Current + (1 | ch_name) + (1 | dyad/subj), data = tf_power_alpha, REML = TRUE, control = lmerControl(optimizer = "bobyqa", calc.derivs = TRUE))`
Linear mixed model fit by REML. t-tests use Satterthwaite's method

REML criterion at convergence: -1325

Scaled residuals:

| Min | 1Q | Median | 3Q | Max |
|---|---|---|---|---|
| -3.24 | -0.62 | -0.06 | 0.59 | 3.68 |

Random effects:

| Groups | Term | Std.Dev. |
|---|---|---|
| subj:dyad | (Intercept) | 0.267492 |
| ch_name | (Intercept) | 0.086437 |
| dyad | (Intercept) | 0.179752 |
| Residual | | 0.174597 |

Number of obs: 2595, groups: subj:dyad, 38; $ch_name, 25; dyad, 20$.

Fixed effects:

| | Estimate | Std. Error | df | t value | Pr(>\|t\|) | |
|---|---|---|---|---|---|---|
| (Intercept) | −8.8 | 0.094 | 32 | −93 | 1.7e−40 | *** |
| role[Commander] | 0.19 | 0.052 | 20 | 3.6 | 0.0019 | ** |
| session[Low] | 0.031 | 0.0051 | 2.5e+03 | 6.2 | 6e−10 | *** |
| session[Med] | −0.034 | 0.005 | 2.5e+03 | −6.7 | 3e−11 | *** |
| study[Armoured] | −0.046 | 0.06 | 18 | −0.76 | 0.46 | |
| $Months_Current$ | −0.0019 | 0.0017 | 31 | −1.2 | 0.26 | |
| role[Commander]:session[Low] | −0.068 | 0.0051 | 2.5e+03 | −13 | 6.3e−40 | *** |
| role[Commander]:session[Med] | 0.034 | 0.005 | 2.5e+03 | 6.7 | 2.9e−11 | *** |
| role[Commander]:study[Armoured] | −0.074 | 0.044 | 17 | −1.7 | 0.11 | |
| session[Low]:study[Armoured] | 0.021 | 0.0051 | 2.5e+03 | 4.2 | 2.9e−05 | *** |
| session[Med]:study[Armoured] | 0.044 | 0.005 | 2.5e+03 | 8.8 | 2.2e−18 | *** |
| role[Commander]:session[Low]:study[Armoured] | −0.02 | 0.0051 | 2.5e+03 | −4 | 6.6e−05 | *** |
| role[Commander]:session[Med]:study[Armoured] | 0.031 | 0.005 | 2.5e+03 | 6.1 | 1.3e−09 | *** |

Table S3: Summary of model produced by the call `lmer(formula = individual_rating ~ power * role * session + Months_Current + (1 | dyad/subj), data = dome_theta, REML = TRUE, control = lmerControl(optimizer = "bobyqa", calc.derivs = TRUE))`
Linear mixed model fit by REML. t-tests use Satterthwaite's method

REML criterion at convergence: 2629

Scaled residuals:

| Min | 1Q | Median | 3Q | Max |
|---|---|---|---|---|
| -2.75 | -0.7 | -0.02 | 0.77 | 3.09 |

Random effects:

| Groups | Term | Std.Dev. |
|---|---|---|
| subj:dyad | (Intercept) | 0.34027 |
| dyad | (Intercept) | 1.03068 |
| Residual | | 0.55880 |

Number of obs: 1486, groups: subj:dyad, 20; dyad, 10.

Fixed effects:

| | Estimate | Std. Error | df | t value | Pr(>\|t\|) | |
|---|---|---|---|---|---|---|
| (Intercept) | 9.9 | 0.58 | 66 | 17 | 6.3e−26 | *** |
| power | 0.33 | 0.057 | 1.3e+03 | 5.8 | 9.1e−09 | *** |
| role[Commander] | −0.53 | 0.44 | 4.1e+02 | −1.2 | 0.23 | |
| session.L | −1.7 | 0.38 | 1.5e+03 | −4.4 | 1.3e−05 | *** |
| session.Q | 0.23 | 0.47 | 1.5e+03 | 0.48 | 0.63 | |
| Months$_{Current}$ | 0.0015 | 0.0051 | 8.7 | 0.29 | 0.78 | |
| power:role[Commander] | −0.068 | 0.057 | 9.7e+02 | −1.2 | 0.23 | |
| power:session.L | −0.19 | 0.051 | 1.5e+03 | −3.7 | 0.00021 | *** |
| power:session.Q | 0.018 | 0.061 | 1.5e+03 | 0.3 | 0.77 | |
| role[Commander]:session.L | 3.1 | 0.38 | 1.5e+03 | 8 | 2.6e−15 | *** |
| role[Commander]:session.Q | −0.14 | 0.47 | 1.5e+03 | −0.3 | 0.76 | |
| power:role[Commander]:session.L | 0.42 | 0.051 | 1.5e+03 | 8.2 | 5.3e−16 | *** |
| power:role[Commander]:session.Q | 0.007 | 0.061 | 1.5e+03 | 0.11 | 0.91 | |

Table S4: Summary of model produced by the call `lmer(formula = individual_rating ~ power * role * session + Months_Current + (1 | dyad/subj), data = dome_alpha, REML = TRUE, control = lmerControl(optimizer = "bobyqa", calc.derivs = TRUE))`
Linear mixed model fit by REML. t-tests use Satterthwaite's method

REML criterion at convergence: 2694

Scaled residuals:

| Min | 1Q | Median | 3Q | Max |
|---|---|---|---|---|
| -2.87 | -0.72 | 0 | 0.6 | 2.85 |

Random effects:

| Groups | Term | Std.Dev. |
|---|---|---|
| subj:dyad | (Intercept) | 0.37189 |
| dyad | (Intercept) | 1.02906 |
| Residual | | 0.57578 |

Number of obs: 1474, groups: subj:dyad, 20; dyad, 10.

Fixed effects:

| | Estimate | Std. Error | df | t value | Pr(>\|t\|) | |
|---|---|---|---|---|---|---|
| (Intercept) | 9.1 | 0.78 | 1.8e+02 | 12 | 9.5e−24 | *** |
| power | 0.2 | 0.078 | 1.2e+03 | 2.6 | 0.01 | * |
| role[Commander] | −0.21 | 0.69 | 6.7e+02 | −0.3 | 0.76 | |
| session.L | −1.4 | 0.66 | 1.5e+03 | −2.2 | 0.028 | * |
| session.Q | −1.2 | 0.71 | 1.5e+03 | −1.6 | 0.1 | |
| Months$_{Current}$ | 0.0027 | 0.0056 | 8.9 | 0.48 | 0.64 | |
| power:role[Commander] | −0.024 | 0.077 | 1e+03 | −0.31 | 0.76 | |
| power:session.L | −0.14 | 0.074 | 1.5e+03 | −1.9 | 0.059 | . |
| power:session.Q | −0.13 | 0.079 | 1.5e+03 | −1.7 | 0.092 | . |
| role[Commander]:session.L | 3 | 0.66 | 1.5e+03 | 4.5 | 6.8e−06 | *** |
| role[Commander]:session.Q | 3.1 | 0.71 | 1.5e+03 | 4.3 | 1.6e−05 | *** |
| power:role[Commander]:session.L | 0.35 | 0.074 | 1.5e+03 | 4.7 | 2.7e−06 | *** |
| power:role[Commander]:session.Q | 0.37 | 0.079 | 1.5e+03 | 4.6 | 4.3e−06 | *** |

Table S5: Summary of model produced by the call `lmer(formula = Score ~ power * role * session + Months_Current + (1 | dyad/subj), data = Tank_Final_Alpha, REML = TRUE, control = lmerControl(optimizer = "bobyqa", calc.derivs = TRUE))`
Linear mixed model fit by REML. t-tests use Satterthwaite's method

REML criterion at convergence: 3007

Scaled residuals:

| Min | 1Q | Median | 3Q | Max |
|---|---|---|---|---|
| -2.87 | -0.67 | -0.09 | 0.74 | 2.7 |

Random effects:

| Groups | Term | Std.Dev. |
|---|---|---|
| subj:dyad | (Intercept) | 6.6975 |
| dyad | (Intercept) | 28.1459 |
| Residual | | 17.8546 |

Number of obs: 351, groups: subj:dyad, 16; dyad, 9.

Fixed effects:

| | Estimate | Std. Error | df | t value | Pr(>\|t\|) | |
|---|---|---|---|---|---|---|
| (Intercept) | 5.2e+02 | 49 | 2.8e+02 | 11 | 1.9e−22 | *** |
| power | 8.3 | 5.4 | 3.1e+02 | 1.5 | 0.12 | |
| role[Commander] | 2e+02 | 43 | 59 | 4.6 | 2.1e−05 | *** |
| session.L | 80 | 59 | 3.3e+02 | 1.4 | 0.17 | |
| session.Q | 86 | 56 | 3.3e+02 | 1.5 | 0.13 | |
| Months$_{Current}$ | 0.022 | 0.094 | 6 | 0.24 | 0.82 | |
| power:role[Commander] | 22 | 4.8 | 59 | 4.7 | 1.5e−05 | *** |
| power:session.L | 10 | 6.6 | 3.3e+02 | 1.6 | 0.11 | |
| power:session.Q | 8.4 | 6.3 | 3.3e+02 | 1.3 | 0.18 | |
| role[Commander]:session.L | −3.7e+02 | 58 | 3.2e+02 | −6.4 | 4.9e−10 | *** |
| role[Commander]:session.Q | 1.5e+02 | 56 | 3.3e+02 | 2.7 | 0.0071 | ** |
| power:role[Commander]:session.L | −41 | 6.5 | 3.2e+02 | −6.4 | 5.7e−10 | *** |
| power:role[Commander]:session.Q | 17 | 6.3 | 3.3e+02 | 2.7 | 0.007 | ** |

Table S6: Summary of model produced by the call `lmer(formula = Score ~ power * role * session + Months_Current + (1 | dyad/subj), data = Tank_Final_Theta, REML = TRUE, control = lmerControl(optimizer = "bobyqa", calc.derivs = TRUE))`
Linear mixed model fit by REML. t-tests use Satterthwaite's method

REML criterion at convergence: 3069

Scaled residuals:

| Min | 1Q | Median | 3Q | Max |
|---|---|---|---|---|
| -2.93 | -0.63 | -0.05 | 0.61 | 2.99 |

Random effects:

| Groups | Term | Std.Dev. |
|---|---|---|
| subj:dyad | (Intercept) | 3.1451 |
| dyad | (Intercept) | 35.0842 |
| Residual | | 18.7539 |

Number of obs: 354, groups: subj:dyad, 17; dyad, 9.

Fixed effects:

| | Estimate | Std. Error | df | t value | Pr(>\|t\|) | |
|---|---|---|---|---|---|---|
| (Intercept) | 4.7e+02 | 31 | 84 | 15 | 1.4e−25 | *** |
| power | 4.6 | 3.7 | 1.1e+02 | 1.2 | 0.21 | |
| role[Commander] | 27 | 25 | 20 | 1.1 | 0.28 | |
| session.L | 1.9e+02 | 39 | 2.9e+02 | 4.9 | 1.4e−06 | *** |
| session.Q | 0.55 | 37 | 3.2e+02 | 0.015 | 0.99 | |
| $Months_{Current}$ | 0.13 | 0.06 | 5.6 | 2.1 | 0.082 | . |
| power:role[Commander] | 3.9 | 3.1 | 20 | 1.3 | 0.22 | |
| power:session.L | 25 | 4.9 | 3e+02 | 5.2 | 3.5e−07 | *** |
| power:session.Q | −1.2 | 4.6 | 3.2e+02 | −0.27 | 0.79 | |
| role[Commander]:session.L | −1.4e+02 | 38 | 2.2e+02 | −3.6 | 0.00033 | *** |
| role[Commander]:session.Q | −48 | 37 | 3.3e+02 | −1.3 | 0.19 | |
| power:role[Commander]:session.L | −17 | 4.7 | 2.2e+02 | −3.5 | 0.00053 | *** |
| power:role[Commander]:session.Q | −6 | 4.6 | 3.3e+02 | −1.3 | 0.19 | |

Table S7: Summary of model produced by the call `lmer(formula = power ~ role * performer * study * session + Months_Current + (1 | ch_name) + (1 | dyad/subj), data = composite_theta, REML = TRUE, control = lmerControl(optimizer = "bobyqa", calc.derivs = TRUE))`
Linear mixed model fit by REML. t-tests use Satterthwaite's method

REML criterion at convergence: 216

Scaled residuals:

| Min | 1Q | Median | 3Q | Max |
|---|---|---|---|---|
| -4.21 | -0.61 | 0.02 | 0.61 | 4.28 |

Random effects:

| Groups | Term | Std.Dev. |
|---|---|---|
| subj:dyad | (Intercept) | 0.49267 |
| ch_name | (Intercept) | 0.11432 |
| dyad | (Intercept) | 0.14800 |
| Residual | | 0.23089 |

Number of obs: 2483, groups: subj:dyad, 37; ch$_name, 25; dyad, 19$.

Fixed effects:

| | Estimate | Std. Error | df | t value | Pr(>\|t\|) | |
|---|---|---|---|---|---|---|
| (Intercept) | −7.8 | 0.15 | 30 | −52 | 5.6e−31 | *** |
| role[Commander] | 0.18 | 0.099 | 21 | 1.9 | 0.078 | . |
| performer[High] | −0.00096 | 0.008 | 2.4e+03 | −0.12 | 0.9 | |
| study[Armoured] | −0.27 | 0.089 | 15 | −3 | 0.0091 | ** |
| session[Low] | −0.049 | 0.0088 | 2.4e+03 | −5.5 | 4.6e−08 | *** |
| session[Med] | −0.054 | 0.0081 | 2.4e+03 | −6.6 | 4e−11 | *** |
| Months$_Current$ | 0.00045 | 0.0029 | 32 | 0.15 | 0.88 | |
| role[Commander]:performer[High] | 0.014 | 0.008 | 2.4e+03 | 1.8 | 0.072 | . |
| role[Commander]:study[Armoured] | −0.056 | 0.082 | 15 | −0.68 | 0.51 | |
| performer[High]:study[Armoured] | −0.016 | 0.008 | 2.4e+03 | −2 | 0.042 | * |
| role[Commander]:session[Low] | −0.17 | 0.0088 | 2.4e+03 | −19 | 3.4e−77 | *** |
| role[Commander]:session[Med] | 0.072 | 0.0081 | 2.4e+03 | 8.9 | 1.1e−18 | *** |
| performer[High]:session[Low] | 0.087 | 0.0096 | 2.4e+03 | 9.1 | 2.8e−19 | *** |
| performer[High]:session[Med] | −0.043 | 0.0085 | 2.4e+03 | −5 | 5.3e−07 | *** |
| study[Armoured]:session[Low] | −0.0005 | 0.0088 | 2.4e+03 | −0.057 | 0.95 | |
| study[Armoured]:session[Med] | 0.051 | 0.0081 | 2.4e+03 | 6.3 | 2.8e−10 | *** |
| role[Commander]:performer[High]:study[Armoured] | 0.0091 | 0.008 | 2.4e+03 | 1.1 | 0.26 | |
| role[Commander]:performer[High]:session[Low] | 0.13 | 0.0096 | 2.4e+03 | 13 | 6.5e−39 | *** |
| role[Commander]:performer[High]:session[Med] | −0.00087 | 0.0085 | 2.4e+03 | −0.1 | 0.92 | |
| role[Commander]:study[Armoured]:session[Low] | −0.11 | 0.0088 | 2.4e+03 | −12 | 6.9e−33 | *** |
| role[Commander]:study[Armoured]:session[Med] | 0.069 | 0.0081 | 2.4e+03 | 8.5 | 2.3e−17 | *** |
| performer[High]:study[Armoured]:session[Low] | 0.075 | 0.0096 | 2.4e+03 | 7.8 | 6.4e−15 | *** |
| performer[High]:study[Armoured]:session[Med] | −0.047 | 0.0085 | 2.4e+03 | −5.5 | 3.9e−08 | *** |
| role[Commander]:performer[High]:study[Armoured]:session[Low] | 0.13 | 0.0096 | 2.4e+03 | 14 | 2e−42 | *** |
| role[Commander]:performer[High]:study[Armoured]:session[Med] | −0.056 | 0.0085 | 2.4e+03 | −6.5 | 8.3e−11 | *** |

Table S8: Summary of model produced by the call `lmer(formula = power ~ role * performer * study * session + Months_Current + (1 | ch_name) + (1 | dyad/subj), data = composite_alpha, REML = TRUE, control = lmerControl(optimizer = "bobyqa", calc.derivs = TRUE))`

Linear mixed model fit by REML. t-tests use Satterthwaite's method

REML criterion at convergence: -1442

Scaled residuals:

| Min | 1Q | Median | 3Q | Max |
|---|---|---|---|---|
| -3.18 | -0.59 | -0.03 | 0.59 | 3.92 |

Random effects:

| Groups | Term | Std.Dev. |
|---|---|---|
| subj:dyad | (Intercept) | 0.270293 |
| ch_name | (Intercept) | 0.087913 |
| dyad | (Intercept) | 0.190125 |
| Residual | | 0.163479 |

Number of obs: 2420, groups: subj:dyad, 36; ch$_name, 25; dyad, 19$.

Fixed effects:

| | Estimate | Std. Error | df | t value | Pr(>\|t\|) | |
|---|---|---|---|---|---|---|
| (Intercept) | −8.9 | 0.1 | 29 | −88 | 3.5e−37 | *** |
| role[Commander] | 0.1 | 0.057 | 19 | 1.8 | 0.09 | . |
| performer[High] | −0.0011 | 0.0057 | 2.4e+03 | −0.19 | 0.85 | |
| study[Armoured] | −0.062 | 0.064 | 17 | −0.97 | 0.35 | |
| session[Low] | 0.0073 | 0.0063 | 2.4e+03 | 1.1 | 0.25 | |
| session[Med] | −0.04 | 0.0058 | 2.3e+03 | −6.9 | 5.5e−12 | *** |
| Months$_{Current}$ | −0.00017 | 0.0018 | 30 | −0.093 | 0.93 | |
| role[Commander]:performer[High] | 0.035 | 0.0057 | 2.4e+03 | 6.1 | 9.4e−10 | *** |
| role[Commander]:study[Armoured] | −0.13 | 0.046 | 16 | −2.8 | 0.012 | * |
| performer[High]:study[Armoured] | −0.005 | 0.0057 | 2.4e+03 | −0.89 | 0.37 | |
| role[Commander]:session[Low] | −0.11 | 0.0063 | 2.4e+03 | −18 | 3e−67 | *** |
| role[Commander]:session[Med] | 0.058 | 0.0058 | 2.3e+03 | 10 | 1.9e−23 | *** |
| performer[High]:session[Low] | 0.029 | 0.0069 | 2.4e+03 | 4.2 | 2.4e−05 | *** |
| performer[High]:session[Med] | −0.031 | 0.0061 | 2.3e+03 | −5.1 | 4.4e−07 | *** |
| study[Armoured]:session[Low] | −0.0048 | 0.0063 | 2.3e+03 | −0.76 | 0.45 | |
| study[Armoured]:session[Med] | 0.037 | 0.0058 | 2.3e+03 | 6.3 | 2.7e−10 | *** |
| role[Commander]:performer[High]:study[Armoured] | 0.028 | 0.0057 | 2.4e+03 | 4.9 | 1.2e−06 | *** |
| role[Commander]:performer[High]:session[Low] | 0.076 | 0.0069 | 2.4e+03 | 11 | 1.8e−27 | *** |
| role[Commander]:performer[High]:session[Med] | −0.011 | 0.0061 | 2.3e+03 | −1.8 | 0.078 | . |
| role[Commander]:study[Armoured]:session[Low] | −0.068 | 0.0063 | 2.3e+03 | −11 | 2.4e−26 | *** |
| role[Commander]:study[Armoured]:session[Med] | 0.051 | 0.0058 | 2.3e+03 | 8.8 | 3e−18 | *** |
| performer[High]:study[Armoured]:session[Low] | 0.038 | 0.0069 | 2.4e+03 | 5.6 | 3.1e−08 | *** |
| performer[High]:study[Armoured]:session[Med] | −0.029 | 0.0061 | 2.3e+03 | −4.7 | 2.9e−06 | *** |
| role[Commander]:performer[High]:study[Armoured]:session[Low] | 0.11 | 0.0069 | 2.4e+03 | 16 | 2.2e−53 | *** |
| role[Commander]:performer[High]:study[Armoured]:session[Med] | −0.039 | 0.0061 | 2.3e+03 | −6.4 | 2.1e−10 | *** |

**Table S9:** Summary of effects of model examining theta across role, session and study

| | Chisq | Df | Pr(>Chisq) |
|---|---|---|---|
| role | 10.8641542 | 1 | 0.0009804 |
| session | 117.2068625 | 2 | 0.0000000 |
| study | 6.4677887 | 1 | 0.0109847 |
| Months_Current | 0.1915799 | 1 | 0.6616049 |
| role:session | 237.6790886 | 2 | 0.0000000 |
| role:study | 0.0141432 | 1 | 0.9053347 |
| session:study | 191.4389080 | 2 | 0.0000000 |
| role:session:study | 88.7225461 | 2 | 0.0000000 |

**Table S10:** Summary of effects of model examining alpha across role, session and study

| | Chisq | Df | Pr(>Chisq) |
|---|---|---|---|
| role | 12.9050171 | 1 | 0.0003277 |
| session | 93.7484328 | 2 | 0.0000000 |
| study | 0.5474413 | 1 | 0.4593651 |
| Months_Current | 1.3230171 | 1 | 0.2500513 |
| role:session | 165.2905681 | 2 | 0.0000000 |
| role:study | 2.8583450 | 1 | 0.0909010 |
| session:study | 172.6115801 | 2 | 0.0000000 |
| role:session:study | 38.4953238 | 2 | 0.0000000 |

**Table S11:** Summary of effects of theta model on behaviour at GBAD

| | Chisq | Df | Pr(>Chisq) |
|---|---|---|---|
| power | 29.8304947 | 1 | 0.0000000 |
| role | 0.0017186 | 1 | 0.9669327 |
| session | 46.4531177 | 2 | 0.0000000 |
| Months_Current | 0.0851113 | 1 | 0.7704867 |
| power:role | 0.2825403 | 1 | 0.5950410 |
| power:session | 34.3994751 | 2 | 0.0000000 |
| role:session | 44.3711766 | 2 | 0.0000000 |
| power:role:session | 68.1091969 | 2 | 0.0000000 |

**Table S12:** Summary of effects of alpha model on behaviour at GBAD

| | Chisq | Df | Pr(>Chisq) |
|---|---|---|---|
| power | 8.0346420 | 1 | 0.0045891 |
| role | 0.0251982 | 1 | 0.8738741 |
| session | 39.7234743 | 2 | 0.0000000 |
| Months_Current | 0.2314303 | 1 | 0.6304653 |
| power:role | 0.8235653 | 1 | 0.3641398 |
| power:session | 1.1520049 | 2 | 0.5621411 |
| role:session | 56.9775739 | 2 | 0.0000000 |
| power:role:session | 39.3305173 | 2 | 0.0000000 |

**Table S13:** Summary of effects of alpha model on behaviour at Armoured

| | Chisq | Df | Pr(>Chisq) |
|---|---|---|---|
| power | 1.2749098 | 1 | 0.2588476 |
| role | 0.6653663 | 1 | 0.4146718 |
| session | 91.1723292 | 2 | 0.0000000 |
| Months_Current | 0.0576444 | 1 | 0.8102585 |
| power:role | 28.4933789 | 1 | 0.0000001 |
| power:session | 18.8508083 | 2 | 0.0000806 |
| role:session | 1.0706166 | 2 | 0.5854887 |
| power:role:session | 44.7294316 | 2 | 0.0000000 |

**Table S14:** Summary of effects of theta model on behaviour at Armoured

| | Chisq | Df | Pr(>Chisq) |
|---|---|---|---|
| power | 3.234076 | 1 | 0.0721210 |
| role | 3.545752 | 1 | 0.0596981 |
| session | 66.453656 | 2 | 0.0000000 |
| Months_Current | 4.468449 | 1 | 0.0345263 |
| power:role | 2.499945 | 1 | 0.1138502 |
| power:session | 41.185424 | 2 | 0.0000000 |
| role:session | 7.223879 | 2 | 0.0269994 |
| power:role:session | 14.594387 | 2 | 0.0006774 |

**Table S15:** Summary of GAMM for alpha power predicting behaviour at GBAD

| *Predictors* | | *Estimate* | *S.E.* | *t* | *p* |
|---|---|---|---|---|---|
| Intercept | | 7.38 | 0.11 | 65.00 | **<.001** |
| Role[Commander] | | -0.33 | 0.11 | -2.92 | **.004** |
| *Smoothers* | | *e.d.f.* | *d.f.* | *F* | *p* |
| ti(power) | | 9.429 | 4 | 4.12 | **<.001** |
| ti(power):RoleCommander | | 3.542 | 4 | 11.05 | **<.001** |
| ti(power):RoleGunner | | 5.459 | 4 | 0.00 | .45 |
| s(channel) | | 5.100 | 7 | 0.00 | .99 |
| Adjusted $R^2$ | .28 | | | | |
| Observations | 160 | | | | |

*Note.* For GAMMS, predictors and smoothers are treatment coded, with commanders set as the reference level for Role. e.d.f. = estimated degrees of freedom (associated *p* values are only approximate); ti() = tensor product interaction term; *s*() = by-channel factor smooth.

**Table S16:** Summary of GAMM for theta power predicting behaviour at GBAD

| *Predictors* | | *Estimate* | *S.E.* | *t* | *p* |
|---|---|---|---|---|---|
| Intercept | | 7.29 | 0.13 | 54.22 | **<.001** |
| Role[Commander] | | -0.12 | 0.13 | -0.93 | .35 |
| *Smoothers* | | *e.d.f.* | *d.f.* | *F* | *p* |
| ti(power) | | 1.231 | 4 | 0.00 | **.02** |
| ti(power):RoleCommander | | 2.702 | 4 | 4.71 | **<.001** |
| ti(power):RoleGunner | | 1.647 | 4 | 1.38 | **.03** |
| s(channel) | | 6.434 | 5 | 0.00 | .99 |
| Adjusted $R^2$ | .17 | | | | |
| Observations | 120 | | | | |

**Table S17:** Summary of GAMM for aperiodic slope predicting behaviour at GBAD

| *Predictors* | | *Estimate* | *S.E.* | *t* | *p* |
|---|---|---|---|---|---|
| Intercept | | 7.46 | 0.10 | 68.99 | **<.001** |
| Role[Commander] | | -0.09 | 0.10 | -0.83 | .40 |
| *Smoothers* | | *e.d.f.* | *d.f.* | *F* | *p* |
| ti(slope) | | 7.558 | 4 | 0.00 | .06 |
| ti(slope):RoleCommander | | 1.583 | 4 | 0.87 | .10 |
| ti(slope):RoleGunner | | 2.216 | 4 | 6.12 | **<.001** |
| s(channel) | | 6.427 | 7 | 0.00 | .99 |
| Adjusted $R^2$ | .15 | | | | |
| Observations | 160 | | | | |

**Table S18:** Summary of GAMM for aperiodic intercept predicting behaviour at GBAD

| *Predictors* | | *Estimate* | *S.E.* | *t* | *p* |
|---|---|---|---|---|---|
| Intercept | | 7.38 | 0.10 | 68.64 | **<.001** |
| Role[Commander] | | -0.09 | 0.10 | -0.09 | .92 |
| *Smoothers* | | *e.d.f.* | *d.f.* | *F* | *p* |
| ti(intercept) | | 9.324 | 4 | 3.44 | **<.001** |
| ti(intercept):RoleCommander | | 2.033 | 4 | 4.28 | **<.001** |
| ti(intercept):RoleGunner | | 1.504 | 4 | 0.87 | .08 |
| s(channel) | | 5.760 | 7 | 0.00 | .99 |
| Adjusted $R^2$ | .17 | | | | |
| Observations | 160 | | | | |

**Table S19:** Summary of general linear model for behaviour predicting behaviour at GBAD

| Predictor | *b* | *b* 95% CI [LL, UL] | $sr^2$ | $sr^2$ 95% CI [LL, UL] | Fit |
|---|---|---|---|---|---|
| (Intercept) | 1.11 | [-4.06, 6.28] | | | |
| individual_rating_Low | 0.83* | [0.16, 1.50] | .29 | [-.04, .63] | |
| role[Commander] | -2.46 | [-7.63, 2.71] | .04 | [-.11, .19] | |
| individual_rating_Low:role[Commander] | 0.31 | [-0.37, 0.98] | .04 | [-.10, .18] | |
| | | | | | $R^2$ = .31 |

*Note.* For general linear models, a significant *b*-weight indicates the semi-partial correlation is also significant. *b* represents unstandardized regression weights. $sr^2$ represents the semi-partial correlation squared. *LL* and *UL* indicate the lower and upper limits of a confidence interval, respectively. * indicates p < .05. ** indicates p < .01.

**Table S20:** Summary of general linear model for IAF predicting behaviour at GBAD

| Predictor | *b* | *b* 95% CI [LL, UL] | $sr^2$ | $sr^2$ 95% CI [LL, UL] | Fit |
|---|---|---|---|---|---|
| (Intercept) | 17.68* | [4.13, 31.23] | | | |
| cog | -1.02 | [-2.38, 0.34] | .14 | [-.14, .42] | |
| role[Commander] | -4.92 | [-18.47,8.63] | .03 | [-.11, .18] | |
| cog:role[Commander] | 0.47 | [-0.89, 1.83] | .03 | [-.11, .17] | |
| | | | | | $R^2$ = .17 |

**Table S21:** Summary of GAMM for alpha power predicting behaviour at Armoured

| *Predictors* | *Estimate* | *S.E.* | *t* | *p* |
|---|---|---|---|---|
| Intercept | 441.25 | 2.65 | 165.98 | **<.001** |
| Role[Commander] | -6.55 | 2.66 | -2.46 | **.01** |
| *Smoothers* | *e.d.f.* | *d.f.* | *F* | *p* |
| ti(power) | 1.493 | 4 | 2.38 | **.002** |
| ti(power):RoleCommander | 2.463 | 4 | 4.22 | **<.001** |
| ti(power):RoleGunner | 5.710 | 4 | 0.00 | .10 |
| s(channel) | 2.707 | 7 | 0.00 | .99 |
| Adjusted $R^2$ | .26 | | | |
| Observations | 80 | | | |

**Table S22:** Summary of GAMM for theta power predicting behaviour at Armoured

| *Predictors* | | *Estimate* | *S.E.* | *t* | *p* |
|---|---|---|---|---|---|
| Intercept | | 441.51 | 2.84 | 155.51 | **<.001** |
| Role[Commander] | | -7.09 | 2.87 | -2.46 | **.01** |
| *Smoothers* | | *e.d.f.* | *d.f.* | *F* | *p* |
| ti(power) | | 8.598 | 4 | 1.53 | **.006** |
| ti(power):RoleCommander | | 2.517 | 4 | 5.61 | **<.001** |
| ti(power):RoleGunner | | 8.686 | 4 | 0.57 | .09 |
| s(channel) | | 5.255 | 5 | 0.00 | .84 |
| Adjusted $R^2$ | .42 | | | | |
| Observations | 60 | | | | |

**Table S23:** Summary of GAMM for aperiodic intercept predicting behaviour at Armoured

| *Predictors* | | *Estimate* | *S.E.* | *t* | *p* |
|---|---|---|---|---|---|
| Intercept | | 442.40 | 2.36 | 187.01 | **<.001** |
| Role[Commander] | | -6.73 | 2.37 | -2.83 | **.005** |
| *Smoothers* | | *e.d.f.* | *d.f.* | *F* | *p* |
| ti(intercept) | | 9.386 | 4 | 3.81 | **.006** |
| ti(intercept):RoleCommander | | 6.944 | 4 | 0.32 | .15 |
| ti(intercept):RoleGunner | | 5.582 | 4 | 0.00 | .87 |
| s(channel) | | 2.862 | 7 | 0.00 | .91 |
| Adjusted $R^2$ | .24 | | | | |
| Observations | 96 | | | | |

**Table S24:** Summary of GAMM for aperiodic slope predicting behaviour at Armoured

| *Predictors* | | *Estimate* | *S.E.* | *t* | *p* |
|---|---|---|---|---|---|
| Intercept | | 442.25 | 2.05 | 215.38 | **<.001** |
| Role[Commander] | | -7.65 | 2.15 | -3.54 | **<.001** |
| *Smoothers* | | *e.d.f.* | *d.f.* | *F* | *p* |
| ti(slope) | | 2.983 | 4 | 15.61 | **.001** |
| ti(slope):RoleCommander | | 3.437 | 4 | 0.00 | .66 |
| ti(slope):RoleGunner | | 8.256 | 4 | 0.00 | .38 |
| s(channel) | | 1.356 | 7 | 0.00 | .62 |
| Adjusted $R^2$ | .42 | | | | |
| Observations | 96 | | | | |

**Table S25:** Summary of general linear model for behaviour predicting behaviour at Armoured

| Predictor | *b* | *b* 95% CI [LL, UL] | $sr^2$ | $sr^2$ 95% CI [LL, UL] | Fit |
|---|---|---|---|---|---|
| (Intercept) | 412.26** | [232.58, 591.93] | | | |
| Score_1 | 0.07 | [-0.33, 0.47] | .01 | [-.09, .11] | |
| role[Commander] | -4.30 | [-183.97, 175.37] | .00 | [-.01, .01] | |
| Score_1:role[Commander] | 0.02 | [-0.38, 0.41] | .00 | [-.02, .02] | |
| | | | | | $R^2$ = .01 |

**Table S26:** Summary of general linear model for IAF predicting behaviour at Armoured

| Predictor | *b* | *b* 95% CI [LL, UL] | $sr^2$ | $sr^2$ 95% CI [LL, UL] | Fit |
|---|---|---|---|---|---|
| (Intercept) | 1057.63** | [672.82, 1442.44] | | | |
| cog | -61.89** | [-101.01, -22.76] | .44 | [.08, .79] | |
| role[Commander] | 234.63 | [-150.18, 619.44] | .07 | [-.10, .23] | |
| cog:role[Commander] | -24.22 | [-63.35, 14.91] | .07 | [-.10, .24] | |
| | | | | | $R^2$ = .55* |

**Table S27.** Summary of the mixed-effects beta regression model.

Family: beta (logit)

Formula: variance ~ study * role * session + (1 | dyad/subj)

| AIC | BIC | logLik | deviance | df.resid |
|---|---|---|---|---|
| -416.5 | -376.3 | 223.3 | -446.5 | 93 |

Random effects:

Conditional model

| Groups | Name | Variance | Std.Dev. |
|---|---|---|---|
| subj:dyad | (Intercept) | 0.007 | 0.08 |
| dyad | (Intercept) | 0.0003 | 0.02 |

Number of obs: 108, groups: subj:dyad, 39; dyad, 20

Dispersion parameter for beta family (): 247

Conditional model:

| | Estimate | Std.Error | z value | p |
|---|---|---|---|---|
| Intercept | -1.15 | 0.02 | -55.69 | <.001 |
| study[Armoured] | 0.02 | 0.02 | 0.94 | 0.35 |
| role[Commander] | -0.01 | 0.02 | -0.81 | 0.41 |
| session[Low] | 0.03 | 0.02 | 1.89 | 0.06 |
| session[Med] | 0.02 | 0.02 | 1.25 | 0.21 |
| study[Armoured]:role[Commander] | 0.01 | 0.02 | 0.81 | 0.41 |
| study[Armoured]:session[Low] | -0.006 | 0.02 | -0.31 | 0.75 |
| study[Armoured]:session[Med] | -0.005 | 0.02 | -0.27 | 0.78 |
| role[Commander]:session[Low] | 0.02 | 0.02 | 0.88 | 0.38 |
| role[Commander]:session[Med] | 0.003 | 0.02 | 0.17 | 0.86 |
| study[Armoured]:role[Commander]:session[Low] | -0.01 | 0.02 | -0.67 | 0.50 |
| study[Armoured]:role[Commander]:session[Med] | -0.002 | 0.02 | -0.11 | 0.91 |

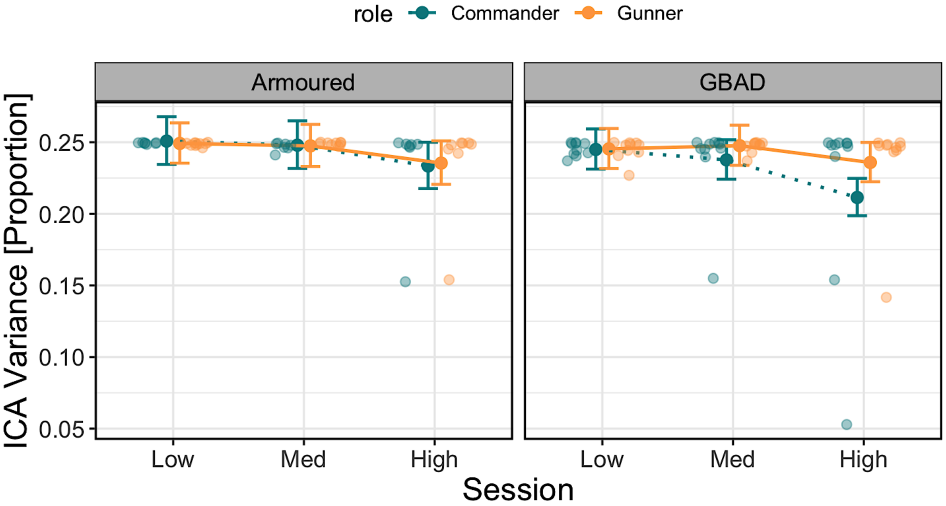

**Figure S1. Modelled effects of the proportion of variance explained by EOG components from the independent component analysis.** (A) Variance explained (y-axis; higher values indicate more proportion of variance explained) across scenario one (Low), two (Med) and three (High; x-axis) between Armoured (left) and GBAD (right). Estimates from commanders are represented by the dashed blue line, while estimates from gunners are represented by the solid orange line. Error bars represent the 83% confidence interval. Individual dots represent raw single-subject data points.
